# Levetiracetam prevents Aβ42 production through SV2a-dependent modulation of App processing in Alzheimer’s disease models

**DOI:** 10.1101/2024.10.28.620698

**Authors:** Nalini R. Rao, Olivia DeGulis, Toshihiro Nomura, SeungEun Lee, Timothy J. Hark, Justin C. Dynes, Emily X. Dexter, Maciej Dulewicz, Junyue Ge, Arun Upadhyay, Eugenio F. Fornasiero, Robert Vassar, Jörg Hanrieder, Anis Contractor, Jeffrey N. Savas

## Abstract

In Alzheimer’s disease (AD), amyloid-beta (Aβ) peptides are produced by proteolytic cleavage of the amyloid precursor protein (APP), which can occur during synaptic vesicle (SV) cycling at presynapses. Precisely how amyloidogenic APP processing may impair presynaptic proteostasis and how to therapeutically target this process remains poorly understood. Using *App* knock-in mouse models of early Aβ pathology, we found proteins with hampered degradation accumulate at presynaptic sites. At this mild pathological stage, amyloidogenic processing leads to accumulation of Aβ_42_ inside SVs. To explore if targeting SVs modulates Aβ accumulation, we investigated levetiracetam (Lev), a SV-binding small molecule drug that has shown promise in mitigating AD-related pathologies despite its mechanism of action being unclear. We discovered Lev reduces Aβ_42_ levels by decreasing amyloidogenic processing of APP in a SV2a-dependent manner. Lev corrects SV protein levels and cycling, which results in increased surface localization of APP, where it favors processing via the non-amyloidogenic pathway. Using metabolic stable isotopes and mass spectrometry we confirmed that Lev prevents the production of Aβ_42_ in vivo. In transgenic mice with aggressive pathology, electrophysiological and immunofluorescent microscopy analyses revealed that Lev treatment reduces SV cycling and minimizes synapse loss. Finally, we found that human Down syndrome brains with early Aβ pathology, have elevated levels of presynaptic proteins, confirming a comparable presynaptic deficit in human brains. Taken together, we report a mechanism that highlights the therapeutic potential of Lev to modify the early stages of AD and represent a promising strategy to prevent Aβ_42_ pathology before irreversible damage occurs.

**One Sentence Summary:** We discovered that the SV-binding drug levetiracetam prevents Aβ_42_ production by modulating SV cycling which alters APP localization and thus proteolytic processing, highlighting its therapeutic potential for AD.

## INTRODUCTION

Alzheimer’s disease (AD) is pathologically characterized by extracellular amyloid plaques and intracellular neurofibrillary tangles (NFTs) composed of amyloid beta (Aβ) peptides and hyperphosphorylated tau, respectively (*1–4*). Amyloid pathology accumulates progressively over 10 or more years in a poorly understood prodromal phase before the manifestation of NFTs, neurodegeneration, and the onset of dementia. Current FDA-approved AD therapeutics are highly effective at removing existing amyloid pathology, but do not stop the production of Aβ peptides (*5, 6*). Therefore, development of a strategy preventing Aβ could minimize downstream neuropathology to prevent or delay AD onset.

Aβ peptides can be generated at presynaptic terminals by sequential proteolytic cleavage of the amyloid precursor protein (APP) through the amyloidogenic processing pathway (*7–13*). This process begins with the cleavage of APP by β-secretase (BACE1), which produces β-CTF. Subsequent proteolytic cleavage of β-CTF by γ-secretase results in the release of Aβ peptides from membranes (*14, 15*). Conversely, in the non-amyloidogenic pathway, APP is cleaved by α-secretase, rather than BACE1, and produces α-CTF. The propensity of APP to undergo cleavage through the amyloidogenic or non-amyloidogenic pathway is strongly influenced by its subcellular localization, as BACE1 activity is increased in acidic environments, such as endosomes and synaptic vesicles (SVs) (*14, 16–18*). Consequently, alterations in presynaptic function might critically influence the processing of APP into Aβ peptides.

A growing body of evidence indicates that presynapses represent important sites for the manifestation of AD pathology (*19–24*). We previously discovered an early impairment in protein degradation in App knock-in (*App* KI) mice that leads to increased levels of presynaptic protein which preceded Aβ peptide accumulation (*25*). Consistently, prior studies of human brains with mild cognitive impairment revealed a similar paradoxical increase in the density of presynaptic puncta (*26*). To address presynaptic dysfunction, we and others have explored the use of Levetiracetam (Lev), an atypical anti-epileptic drug (AED) that binds the synaptic vesicle glycoprotein 2A (SV2a) protein at axon terminals (*27–29*). Despite the wide use of Lev for decades to effectively quell seizures in humans, its precise molecular mechanism remains unclear (*30*). Lev has emerged as a potential therapeutic for AD since it mitigates excess synaptic activity and because Aβ can be produced and released through the SV cycle (*8, 9, 13, 31*). Interestingly, in AD animal models and AD patients, Lev treatment reduces plaque pathology, memory deficits, and slows cognitive decline (*27, 28, 32–35*). The therapeutic benefit of Lev is notably not based solely on its ability to reduce excessive synaptic activity, as typical AEDs were equally effective at reducing hyperactivity but did not improve performance on cognitive tasks (*28, 36*). Lev is currently the subject of several clinical trials for AD, but the mechanisms by which it helps reduce AD pathology are still unclear.

In this study, we investigated the therapeutic mechanism of action of Lev to modulate amyloid pathology. First, we followed up on our previous findings that presynaptic proteins have impaired degradation prior to significant Aβ accumulation in *App* KI brains (*25*). We found that proteins with impaired degradation are present at presynaptic sites. Biochemical isolation of SVs revealed that Aβ_42_ is accumulated in the SV lumen. We then determined that Lev modulates APP proteolytic processing by correcting SV protein levels and decreasing SV cycling. As a result, APP is preferentially localized at the plasma membrane, where it is more likely to be processed by the non-amyloidogenic pathway, thereby reducing Aβ_42_ levels. Notably, Lev decreases amyloid pathology *in vivo* by preventing production of Aβ_42_ and minimizes synapse loss. Finally, we performed quantitative proteomic analyses of human Down syndrome (DS) brains and found elevated levels of presynaptic proteins prior to significant Aβ_42_ accumulation. Taken together, our findings document impaired presynaptic protein degradation early in amyloid pathology and reveals the therapeutic mechanism of action for Lev to prevent the production of Aβ_42_ and consequently, downstream irreversible damage.

## RESULTS

### Proteins with impaired degradation accumulate at presynaptic sites during the early stages of Aβ_42_ accumulation

We previously determined that protein turnover is impaired in *App^NL-F/NL-F^*(*NL-F*) and *App^NL-G-F/NL-G-F^* (*NL-G-F*) relative to *App^NL/NL^* (*NL*) knock-in mice of amyloid pathology (*25*). Both *NL-F* and *NL-G-F* mice have elevated levels of presynaptic proteins and excess SVs. Notably, protein turnover was impaired in *NL-F* mice before elevated Aβ_42_ levels or plaques could be detected (*25*). To further our understanding of this proteostasis deficit, we sought to determine where proteins with impaired degradation accumulate in the *NL-F* brain. ELISA analysis of the soluble and insoluble fractions from cortical homogenates confirmed a slight non-significant increase in Aβ_42_ levels in *NL-F* and substantial Aβ_42_ increase in *NL-G-F* compared to *NL* controls (**Fig. 1A-B**). Next, we used a transgenic mouse line expressing a readily degradable GFP* (i.e., *G76V-GFP*) as a sensor to visualize where proteins with hampered degradation accumulate (*37, 38*). In *G76V-GFP/NL-F* mice, we observed significantly increased GFP* intensity in the cortex but not the cerebellum compared to *G76V-GFP* mice **(Fig. 1C-D)**. GFP* also colocalized with ubiquitin puncta but not mitochondria **(Fig. S1A-B)**. To investigate the possibility that Aβ_42_ is present near the GFP* sensor, we utilized immunofluorescence (IF) analysis to quantify the colocalization. GFP* intensity at Aβ_42_ puncta was significantly higher in *G76V-GFP/NL-F* compared to *G76V-GFP* (**Fig. 1E-F, Fig. S1C-D)**. To probe if GFP* accumulates in extracellular plaques, we performed IF analysis of *G76V-GFP/NL-G-F* brain. GFP* does not colocalize with extracellular Aβ_42_ deposits (**Fig. S1E**). Using IF analysis, we investigated the possibility that GFP* is accumulating at or near synapses (i.e., defined as Bassoon and PSD95 positive puncta). This revealed that GFP* intensity is significantly increased at synaptic puncta in *G76V-GFP/NL-F* mice compared to controls **(Fig. 1G-H)**. Furthermore, over 90% of the total GFP* signal colocalizes with synaptic puncta (**Fig. S1F**). To further dissect if GFP* is closer to presynaptic or postsynaptic sites, we performed super resolution microscopy on *G76V-GFP/NL-F* sections. To determine if GFP* is in closer proximity to Bassoon or PSD95 puncta, we compared the peak of each intensity distribution from multiple synapses across biological replicates. The peak-to-peak distance from GFP* to Bassoon was significantly shorter than the distance to PSD95, indicating that GFP* accumulates closer to presynaptic sites **(Fig. 1I-J)**. As Aβ_42_ deficits have previously been reported to differentially affect excitatory versus inhibitory synapses, we tested if GFP* colocalized with VGluT1 or VGAT puncta (*39–41*). In *G76V-GFP/NL-F* brains, GFP* colocalizes to a significantly greater degree to excitatory (VGluT1) rather than inhibitory (VGAT) presynaptic puncta **(Fig. S1G-H)**. Furthermore, biochemical isolation of pre- and postsynaptic fractions using sucrose gradients and WB analysis validated that GFP* predominately present in synaptosomes and the presynaptic fraction but not the postsynaptic fraction (*42, 43*) **(Fig. 1K, Fig. S1I)**. Altogether, in *NL-F* brains with early Aβ_42_ pathology, proteins accumulate at presynaptic sites.

**Figure 1.**
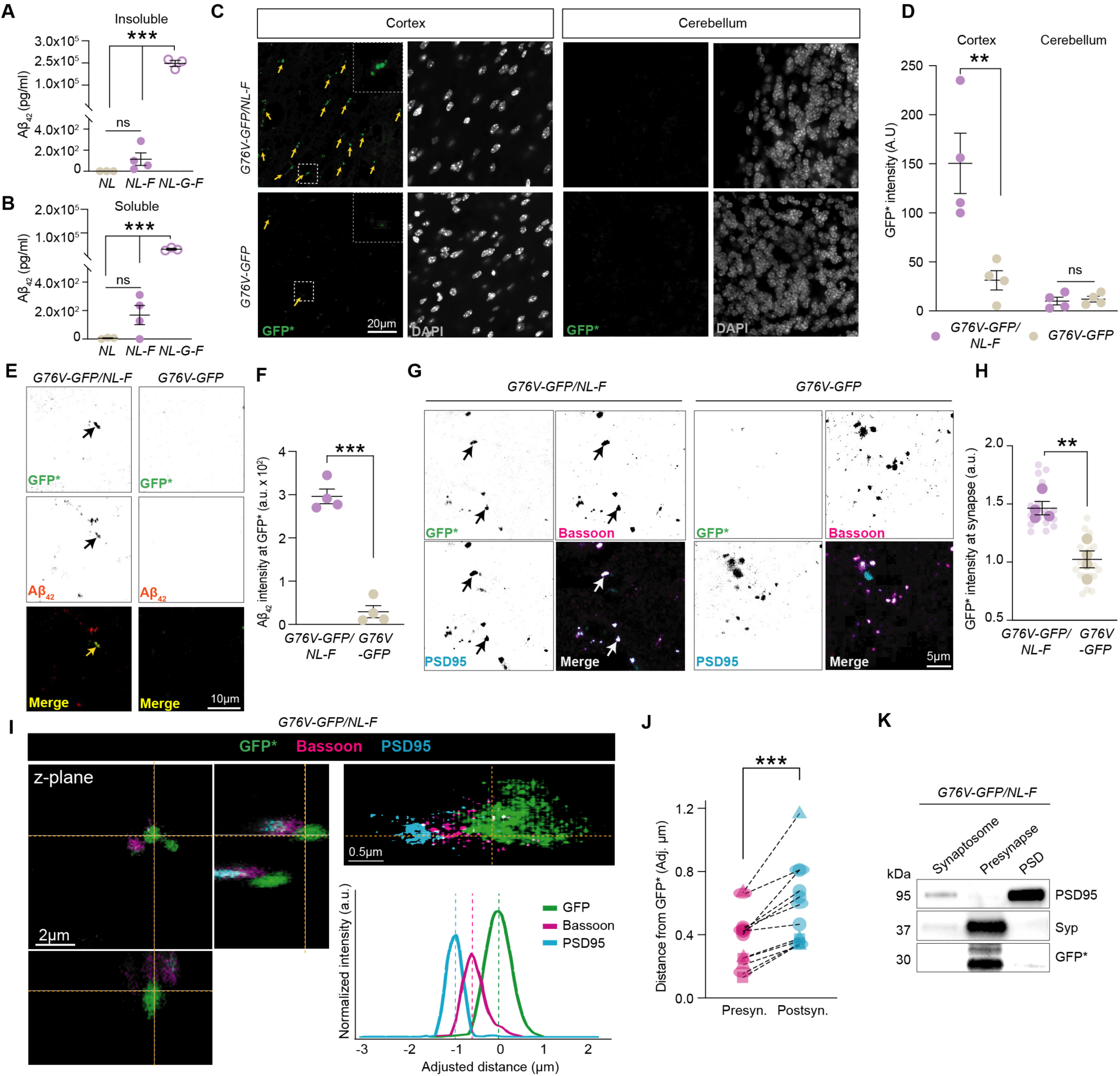
Proteins with impaired degradation accumulate at presynaptic sites during early stages of Aβ_42_ levels. (A) Aβ_42_ levels in the insoluble fraction of 6 months old *NL-F* are slightly elevated based on Aβ_42_ sandwich ELISA compared to *NL-G-F* positive controls and *NL* negative controls. (B) Aβ_42_ levels in the soluble fraction of 6 months old *NL-F* are slightly elevated based on Aβ_42_ sandwich ELISA compared to *NL-G-F* positive controls and *NL* negative controls. (C) Representative IF image of *G76V-GFP/NL-F* mice showing GFP* signal in the cortex, but not the cerebellum, compared to *G76V-GFP* control mice. Scale bar is 20µm. (D) Quantitation of (C) showing that *G76V-GFP/NL-F* mice have significantly increased intensity of GFP* signal compared to *G76V-GFP* control mice in the cortex but not the cerebellum. (E) Representative IF image showing *G76V-GFP/NL-F* mice have Aβ_42_ colocalization at GFP* puncta. Scale bar of 10µm. (F) Quantitation of (E). *G76V-GFP/NL-F* mice have significantly higher Aβ_42_ intensity at GFP* puncta than *G76V-GFP* control mice. Intensity of Aβ_42_ signal at GFP* positive puncta was quantified in cortical areas. (G) Representative IF image showing *G76V-GFP/NL-F* mice have GFP* that colocalizes with synaptic markers (Bassoon and PSD95). Scale bar is 5µm. (H) Quantitation of (G). Intensity of GFP* puncta is significantly higher at synaptic puncta, in *G76V-GFP/NL-F* compared to *G76V-GFP* control mice. GFP* intensity was extracted from puncta positive for Bassoon and PSD95 and normalized to *G76V-GFP*. (I) Representative super resolution microscopy image of *G76V-GFP/NL-F* mice reveals that GFP* is closer to presynaptic puncta. Representative intensity distributions for GFP*, Bassoon, and PSD95 shows that GFP* overlaps with Bassoon and not PSD95. Scale bar is 2µm and 0.5µm. (J) Quantitation of (I). GFP* is significantly closer to presynaptic puncta compared to postsynaptic puncta. Adjusted distances from peak of intensity distribution for Bassoon and PSD95 compared to GFP* were quantified and three synapses per biological replicate were analyzed with a paired t-test. (K) Representative WB showing GFP* is present in the presynaptic but not postsynaptic biochemical fraction from *G76V-GFP/NL-F* from the cortex. All data are mean ± SEM with n = 3-4 mice at 6 months of age. ** = p value < .01; *** = p value < .001; by Student’s t-test (A, B, D, F, H) or paired t-test (J).

### Synaptic vesicles harbor full length App, CTFs, and Aβ_42_

Presynapse function revolves around the SV cycle and represents a highly dynamic cellular process. App can be endocytosed from the plasma membrane (PM) and its topology, together with cleavage sites, within the lumen and SV membrane plays a key role in proteolytic processing (*11*). Amyloidogenic processing of App has been shown to occur at presynapses and Aβ_42_ is released into the extracellular space by SV cycling (*8–10*). First, we purified SVs from *NL-F* and WT mouse brains using synaptosome isolation and size exclusion chromatography then performed EM and proteomics (**Fig. 2A-C, Fig. S2A**). WB analysis showed that PSD95 and Lamp1 were generally absent, while β- and γ-secretases were present in the purified SV material (**Fig. 2D, Fig. S2B**). We next used proteinase K (PK)-based proteolysis to confirm the previously reported orientation of App on SVs. WB analysis using antibodies recognizing luminal or cytosolic Syt1 epitopes, in addition to SV2a and Vamp2, demonstrated that indeed, PK treatment only cleaved the cytosolic domains, while the luminal domain of Syt1 remained physically inaccessible **(Fig. 2E-F, Fig. S2C)**. Consistent with previous reports, we also confirmed that the N- terminus of App is located in the SV lumen, whereas the C- terminus is facing the cytosol in *NL-F* and WT SVs **(Fig. 2G-H, Fig. S2D)**.

**Figure 2.**
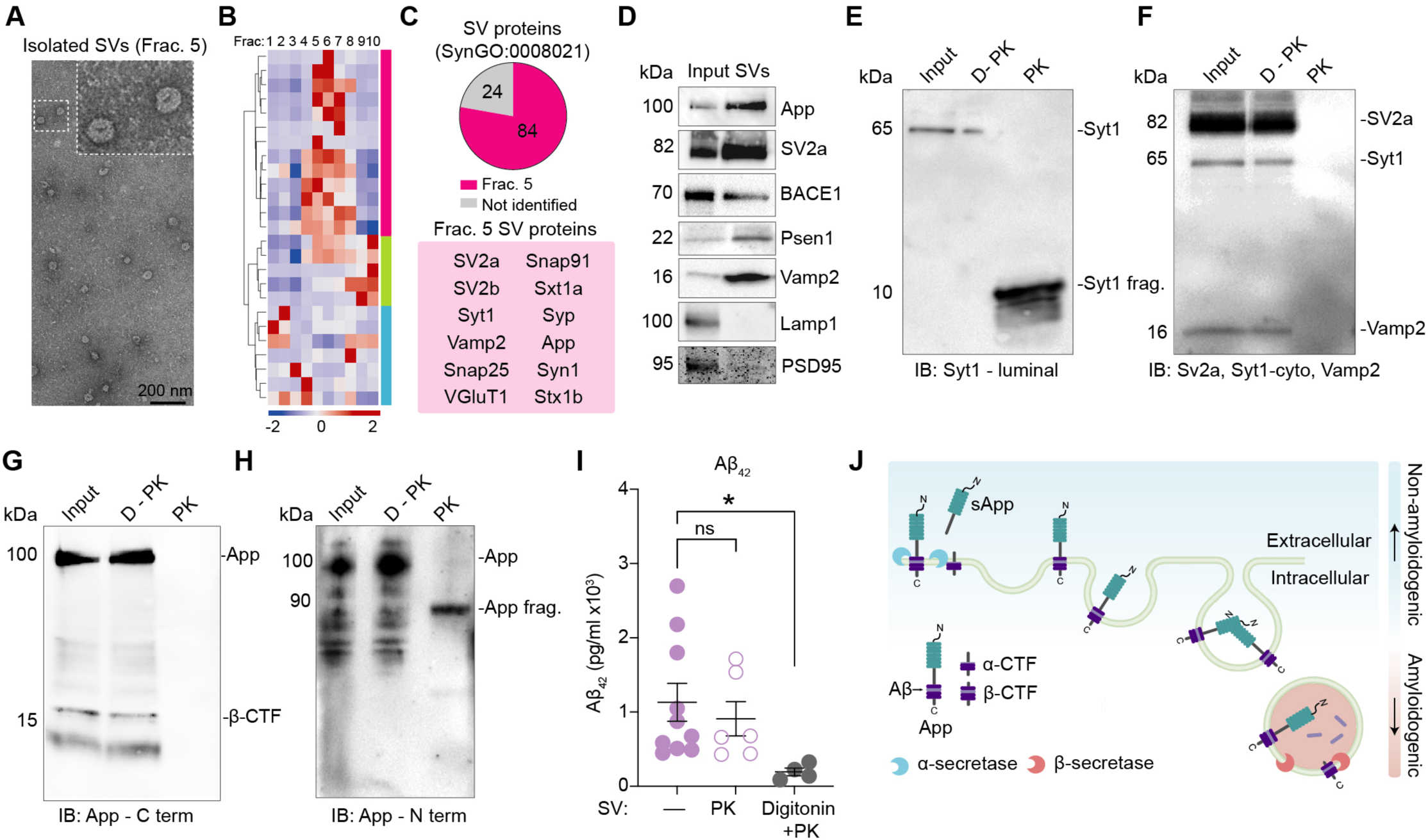
Synaptic vesicles harbor App, CTFs, and Aβ_42_. (A) Representative electron micrograph of Fraction 5 (Frac. 5) depicting abundant SVs from *NL- F* cortical extracts. Scale bar is 200 nm. (B) Heatmap depicting z-score abundance of the proteins identified from MS-based proteomic analysis of the IZON SEC fractions (Frac.1-10). (C) Frac. 5 contains highest levels of many SV proteins and pie chart illustrates that Frac. 5 also contains SV proteins based on SynGO:0008021. A panel of the Frac. 5 proteins identified are shown in the red box. (D) Representative WB analysis of Frac. 5 showing enrichment of SV proteins from *NL-F* cortical homogenates. (E-F) WB analysis of SVs (Input), SVs treated with heat deactivated PK (D-PK), and SVs treated with PK (PK) probed with a luminal Syt1 antibody confirm proteolytic digestion removed of cytoplasmic epitope of Syt1, while leaving the luminal fragment (∼10 kDa) intact. Probing for cytoplasmic epitopes of SV2a, Syt1, and Vamp2 confirm that PK treatment effectively removed cytoplasmic proteins from intact SVs. (G-H) WB analysis of SVs (Input), SVs treated with heat deactivated PK (D-PK), and SVs treated with PK (PK) probed with a C-terminal App antibody shows that no detection of C- terminal App epitope is present after PK treatment, indicating its cytoplasmic orientation. While the N-terminal App antibody shows that the remaining N-terminal App fragment is detected at ∼90 kDa after PK treatment, indicating its orientation is facing the lumen. (I) Aβ_42_ ELISA analysis of *NL-F* SVs treated with PK or detergent (Digitonin) + PK reveals that Aβ_42_ is in the lumen of SVs. (J) Schematic depicting App proteolytic processing pathways. All data are mean ± SEM with n = 4-10 mice at 6 months of age. * = p-value < .05 by ANOVA with Tukey’s multiple comparisons test (I).

Utilizing this PK assay with ELISA as the readout, we found that Aβ_42_ levels in SVs are not affected by PK treatment unless the SV membrane is physically disrupted and made accessible with detergent **(Fig. 2I).** This indicates that Aβ_42_ is predominantly located in the lumen of SVs rather than being present outside of the SVs. Lastly, to investigate whether Aβ_42_ is preferentially associated with specific SVs, we used SV2a-based immunocapture with IF analysis **(Fig. S2E)**. Notably, we found that Aβ_42_ puncta colocalized significantly more with VGluT1 positive SVs than with VGAT SVs **(Fig. S2F-G).** Thus, these results show that the orientation of App within SV membranes favors BACE1 proteolysis and subsequent γ-secretase proteolytic processing generates Aβ_42_ in SVs which may disrupt the SVC (**Fig. 2J**).

### Pharmacological targeting of SVs with levetiracetam modulates APP processing in an SV2a-dependent manner

We next tested if the SV-binding small molecule levetiracetam (Lev) affects Aβ_42_ levels. First, we established an *in vitro* overexpression platform that allows mechanistic examination of Lev’s action on APP processing and Aβ_42_ production. We infected primary rodent neurons with lentiviruses (LVs) expressing human APP or APP with Swedish and Indiana (K670_M671, V717F, APP^Swe/Ind^) mutations (*44*). Neurons overexpressing APP^Swe/Ind^ had increased levels of β-CTF compared to neurons overexpressing full length APP at the same level (**Fig. S3A-C**). We confirmed that APP^Swe/Ind^ neurons generate Aβ_42_ in a manner dependent on BACE1 (C3) and γ-secretase (DAPT) (**Fig. 3A, Fig. S3D**). To confirm this *in vitro* overexpression platform recapitulates our previous finding that presynaptic protein turnover is impaired, we used metabolic pulse-chase labeling in combination with quantitative proteomics (**Fig. S3E-F**) (*25*). This revealed a panel of proteins with hampered turnover in APP^Swe/Ind^ expressing neurons **(Fig. S3G)**. Many of these proteins are associated with the presynapse and were previously found to exhibit hampered turnover in our studies of *NL-F* mice **(Fig. S3H)** (*25*).

**Figure 3.**
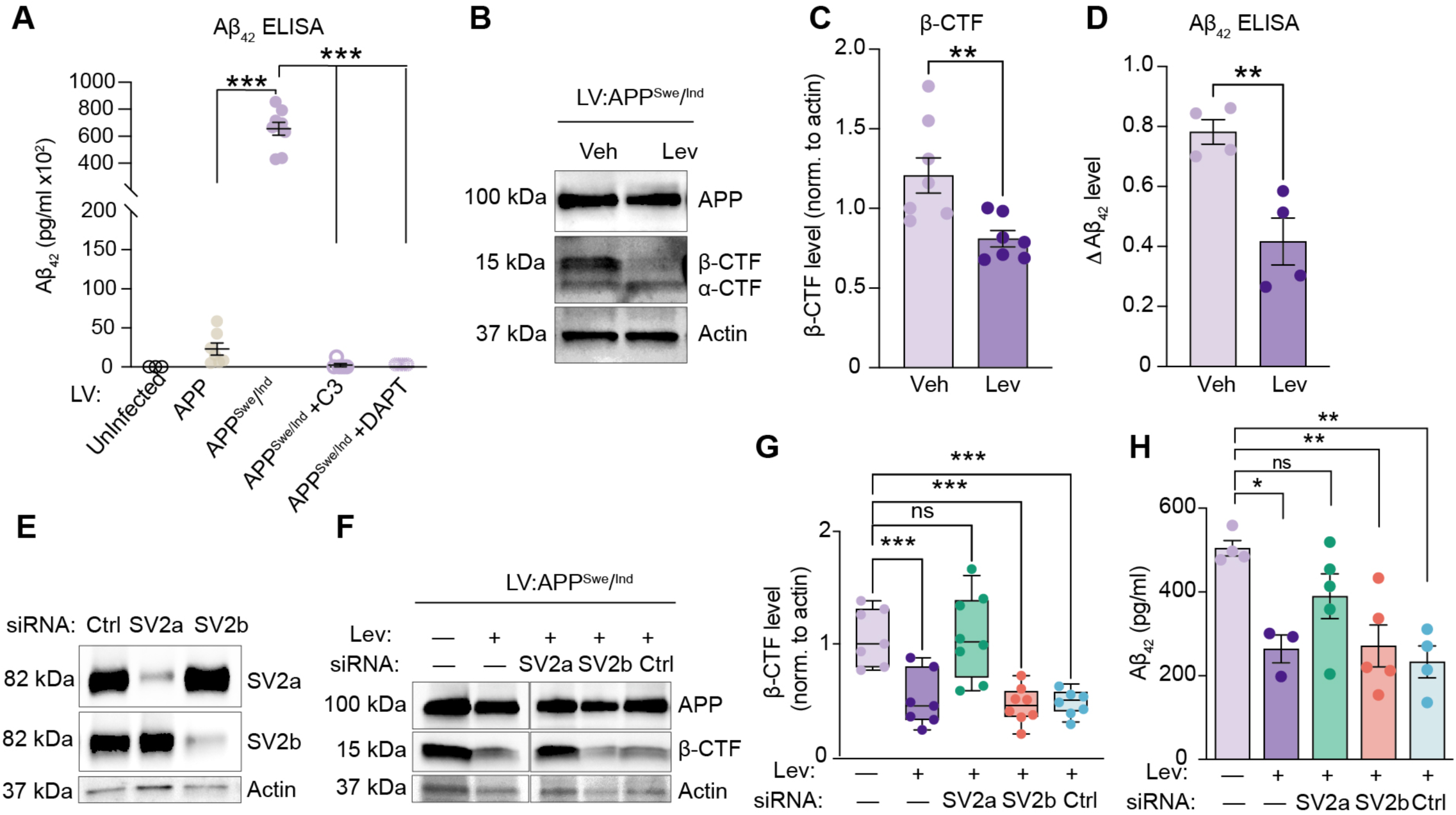
SV-targeting levetiracetam modulates amyloidogenic APP processing in SV2a-dependent manner. (A) ELISA analysis of Aβ_42_ levels from media confirms that APP^Swe/Ind^ neurons produce significantly more Aβ_42_ compared to APP neurons. This production of Aβ_42_ is prevented by treating APP^Swe/Ind^ neurons with β- and γ- secretase inhibition (C3 and DAPT, respectively). (B-C) Representative WB and quantification of APP^Swe/Ind^ neurons treated with 150 µM Lev for 24 hrs shows a significant decrease in β-CTF abundance compared to Veh. WB quantification is normalized to actin. (D) ELISA quantification of Aβ_42_ levels from media of APP^Swe/Ind^ neurons treated with 150 µM Lev for 24 hours shows a significant decrease in Aβ_42_ levels compared to Veh. Media was collected before and after Lev treatment and the change (Δ) in Aβ_42_ levels after Lev is plotted. (E) Representative WB shows effective siRNA-based knock down of SV2a and SV2b compared to non-targeting pool control (Ctrl). (F) Representative WB analysis of APP^Swe/Ind^ neurons treated with Lev or Veh shows that SV2a is required for Lev to reduce β-CTF levels. (G) Quantification of (F). APP^Swe/Ind^ neurons treated with Lev have significantly reduced β-CTF levels unless SV2a is removed. (H) ELISA analysis confirms that SV2a is required for Lev to significantly decrease Aβ_42_ levels in APP^Swe/Ind^ neurons. All data are mean ± SEM with n = 3-8 biological replicates. * = p value < .05; ** = p value < .01; *** = p value < .001; by Student’s t-test for (C, D) or ANOVA with Dunnet’s multiple comparisons test for (A,G, H).

SV2a is a direct binding target of Lev (*30, 45, 46*). However, the molecular mechanism of action underlying Lev’s ability to reduce amyloid pathology, and whether it depends on SV2a, remains unknown. We discovered that APP^Swe/Ind^ expressing neurons incubated with Lev have a robust decrease of β-CTF and Aβ_42_ levels, but not full-length APP levels, compared to vehicle (Veh) (**Fig. 3B-D, Fig. S4A-B**). This suggests that Lev reduces β-CTF and Aβ_42_ levels by modulating APP processing, not APP abundance itself. To address if this reduction requires SV2a, we utilized siRNA knockdown combined with Lev treatment. WB analysis confirmed siRNA-based knock down (KD) of SV2a and SV2b in neurons and that reducing levels of SV2a did not reduce SV2b levels or vice versa **(Fig. 3E)**. Similarly, KD of SV2a or SV2b in absence of Lev treatment did not affect β-CTF levels (**Fig. S4C**). Finally, combination of siRNA-based KD with Lev treatment revealed that SV2a is required for Lev to reduce β-CTF and Aβ_42_ levels **(Fig. 3F-H)**. Taken all together, these results confirm that Lev reduces amyloidogenic processing of APP through SV2a.

### Levetiracetam corrects SV cycling and increases plasma membrane localization of APP

To next investigate how Lev alters the proteome, we performed a tandem mass tag (TMT)-MS experiment on APP^Swe/Ind^ neurons treated with Veh, Lev, or siRNA KD of SV2a + Lev. Importantly, on average there was no global difference in relative protein abundance between the three groups (**Fig. 4A, Fig. S5A, and Table S1**). Performing a Bayesian analysis of variance of the Lev and SV2a KD + Lev groups, each with respect to Veh, revealed that Lev led to substantially more significantly modulated proteins compared to the SV2a KD + Lev (**Fig. 4B, Fig. S5B**). Gene ontology (GO) enrichment analysis of the proteins significantly modulated by Lev showed that proteins associated with membranes and vesicles were overrepresented (**Fig. S5C**). Given the evidence linking Lev and the presynapse, we next extracted the proteins in our dataset associated with synapses using SynGO (n = 588) (*27*). Notably, Lev significantly decreased synaptic protein levels in an SV2a-dependent manner (**Fig. 4C**). We next extracted proteins with a decrease in abundance in the Lev group compared to both the Veh and SV2a KD + Lev groups (n = 203 proteins). GO analysis of these proteins revealed that the most significantly enriched term is Synaptic Vesicle Cycle (GO:0099504) (**Fig. S5D**). This group included Syt1 (an SV2a interactor), Rab5c, Ap1b1, and Snap91, among others (**Fig. 4D**). To investigate the effect of Lev treatment on wild type non-Aβ producing neurons, we performed an additional TMT-based proteomic experiment and found that presynaptic proteins were not significantly altered (**Fig. S5E-F**).

**Figure 4.**
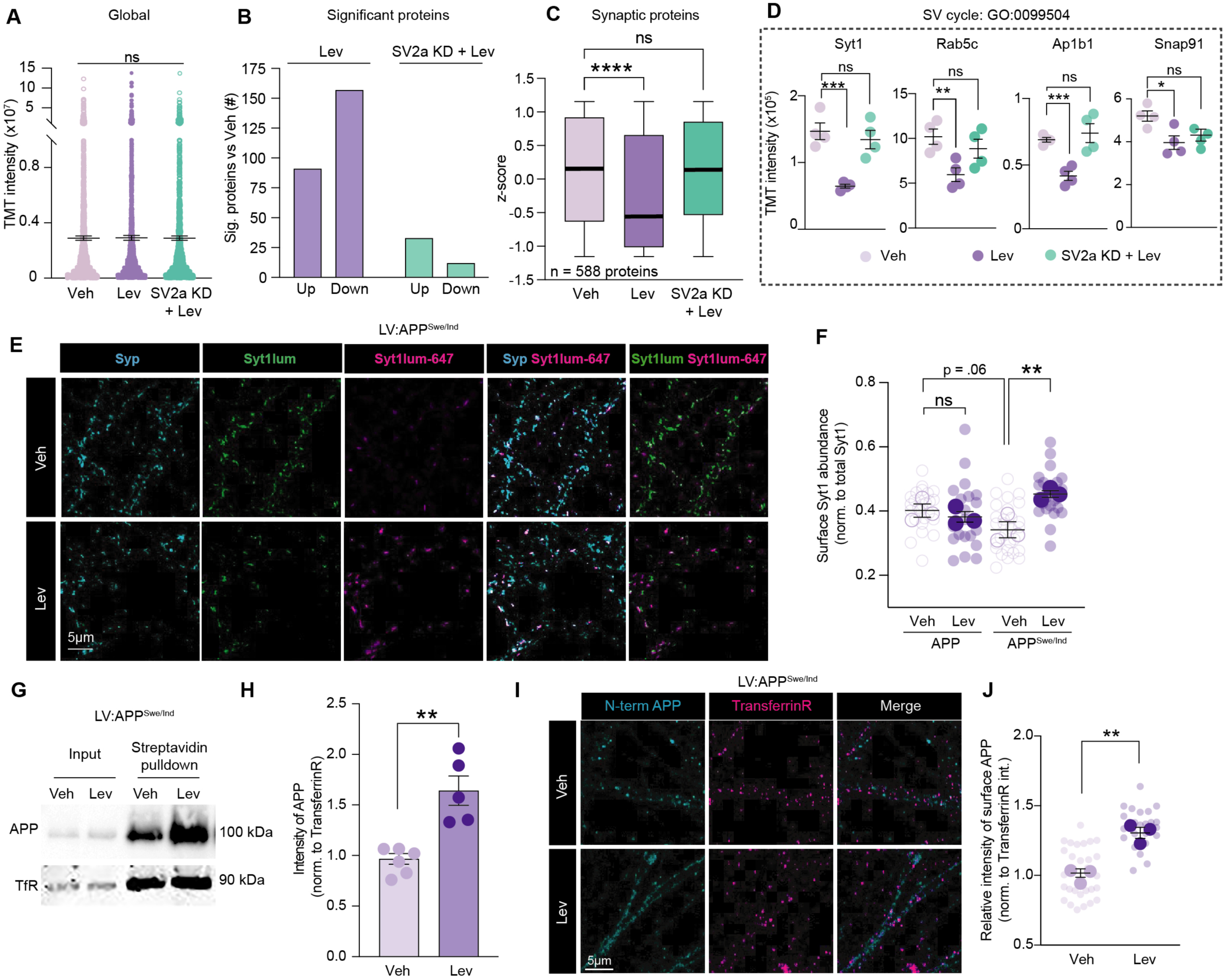
Levetiracetam decreases SV cycling and corrects elevated levels of presynaptic proteins leading to plasma membrane localization of APP. (A) TMT-MS proteomic analysis was performed on APP^Swe/Ind^ expressing neurons treated with Veh, Lev, or SV2a siRNAs with Lev (SV2a KD + Lev). Global TMT reporter ion intensities comparing the proteomes of each group showed no significant average difference. (B) Number of significantly modulated proteins based on Bayesian analysis of variance revealed that Lev treatment had a substantial effect on the proteome that requires SV2a. (C) Average TMT intensities of the proteins classified as synaptic based on SynGO, shows Lev significantly decreases presynaptic protein abundance in an SV2a-dependent manner. (D) A panel of SV Cycle (GO:0099504) proteins displaying significantly decreased relative abundance with Lev treatment in an SV2a-dependent manner. (E) Representative Veh or Lev treated APP^Swe/Ind^ expressing neurons after live-cell incubation with Syt1luminal-647 antibody to visualize the pool of Syt1 on the surface. After live-cell labeling, neurons were fixed, permeabilized, and immunostained for synaptophysin (Syp) and a second Syt1 luminal antibody revealing the internal epitopes. Synaptophysin (cyan), Syt1lum (green), Syt1lum-647 (magenta). Scale bar is 5µm. (F) Quantification of (E). Number of Syt1lum-647 surface puncta relative to total Syt1 puncta (Syt1lum-647 + Syt1lum) from Veh and Lev treated APP and APP^Swe/Ind^ expressing neurons. Lev treatment resulted in a significant increase in surface Syt1 (Syt1lum-647) relative to total Syt1. (G) Representative WB analysis of APP and TransferrinR levels from surface biotin labeling and streptavidin capture of Veh or Lev treated APP^Swe/Ind^ neurons. (H) Quantification of (G). Lev treated APP^Swe/Ind^ expressing neurons have significantly more APP expressed on the surface compared to Veh. Abundance of surface APP was normalized to TransferrinR. (I) Representative Veh or Lev treated APP^Swe/Ind^ expressing neurons after live-cell incubation with an N-terminal APP antibody then fixed and immunostained for TransferrinR without permeabilization. Scale bar is 5µm. (J) Quantification of (I). Lev treatment results in significantly increased surface APP intensity compared to Veh. Intensity of surface APP was normalized to surface TransferrinR intensity. All data are mean ± SEM with n = 3-6 biological replicates. * = p value < .05; ** = p value < .01; *** = p value < .001; **** = p value < .0001 by Student’s t-test for (H, J) or ANOVA with Dunnet or Sidak multiple comparisons test (A, C, D). One-way ANOVA with post-hoc one-sided t-tests were performed (F).

As Lev decreased levels of SV proteins, we next aimed to address how Lev impacts SV exo/endocytosis (i.e. cycling) dynamics. To accomplish this, we performed a live cell surface Syt1-luminal-647 antibody binding assay in WT, APP, and APP^Swe/Ind^ neurons (*47*). SV retrieval during cycling can be measured by comparing the level of surface accessible Syt1-luminal epitopes (*48*). As we found that Syt1 levels were reduced by Lev, we aimed to determine if Lev increased the abundance of surface Syt1 relative to total Syt1. After acute incubation with a Syt1lum-647 antibody, neurons were gently fixed and permeabilized before immunostaining with a second Syt1-lum antibody made in a different species but with the same epitope to detect the remaining unlabeled pool of Syt1. In both APP and APP^Swe/Ind^ expressing neurons, the colocalization of Syt1lum-647 with Synaptophysin was significantly greater compared to the colocalization of Syt1lum-647 with Syt1-lum (**Fig. S5G**). This confirms our paradigm detects two discrete pools of Syt1. Next, we tested how Lev affects SV cycling in APP^Swe/Ind^ expressing neurons and found that treatment significantly increased abundance of surface Syt1 compared to Veh (**Fig. 4E-F)**. Also, Veh treated APP expressing neurons had increased surface Syt1 compared to Veh APP^Swe/Ind^ neurons. Additionally, Lev treatment did not modulate surface Syt1 levels in WT nor APP expressing neurons (**Fig. 4F, Fig. S5H)**.

Non-amyloidogenic APP processing occurs preferentially at the cell surface on the plasma membrane (PM) (*7, 49*). We hypothesized that as Lev alters SV cycling, this would affect the localization of full-length APP. To address this, we first performed live-cell labeling of the surface proteome using biotin to quantify APP PM levels after Lev or Veh treatment in APP^Swe/Ind^ neurons. Streptavidin capture of the biotinylated proteins followed by WB analysis revealed that Lev significantly increased PM APP levels relative to the ubiquitous surface protein transferrin receptor (**Fig. 4G-H**). Next, we performed additional live cell labeling with an APP N-terminal antibody in APP^Swe/Ind^ expressing neurons incubated with Lev or Veh. Consistent with the biochemical experiment, Lev significantly increased total PM APP intensity (**Fig. 4I-J**). These results indicate that Lev treatment corrects SV dynamics, leading to increased levels of full-length APP^Swe/Ind^ on the surface.

### Levetiracetam prevents Aβ_42_ production *in vivo*

Our finding that Lev decreased amyloidogenic processing *in vitro* led us to address whether chronic Lev treatment alters App processing *in vivo* in the *NL-F* model (**Fig. S6A**). Aβ_42_ ELISA analysis of cortical homogenates showed that Lev decreased Aβ_42_ levels (p value = .06), however we noted a clear distinction of two subpopulations by sex (**Fig. S6B**). Lev treated female *NL-F* mice exhibited significantly reduced Aβ_42_ levels compared to Veh, and male *NL-F* mice displayed a similar trend (**Fig. 5A, Fig. S6C**). Next, to investigate how Lev modulated the proteome, we performed TMT-MS proteomics which revealed many proteins associated with SV cycling had significantly reduced levels after Lev treatment (**Fig. S6D-F**). Additionally, TMT-MS confirmed that Lev reduced Aβ peptide levels while App levels remained unchanged (**Fig. 5B)**. Consistently, WB analysis confirmed that Lev did not affect the level of full-length App, but did reduce β-CTF levels (**Fig. 5C-D, Fig. S6G**). Finally, in addition to studying the fragments generated during amyloidogenic processing, we measured sAppα levels, the byproduct of non-amyloidogenic processing. We found that Lev significantly increased sAppα abundance (**Fig. 5E**).

**Figure 5.**
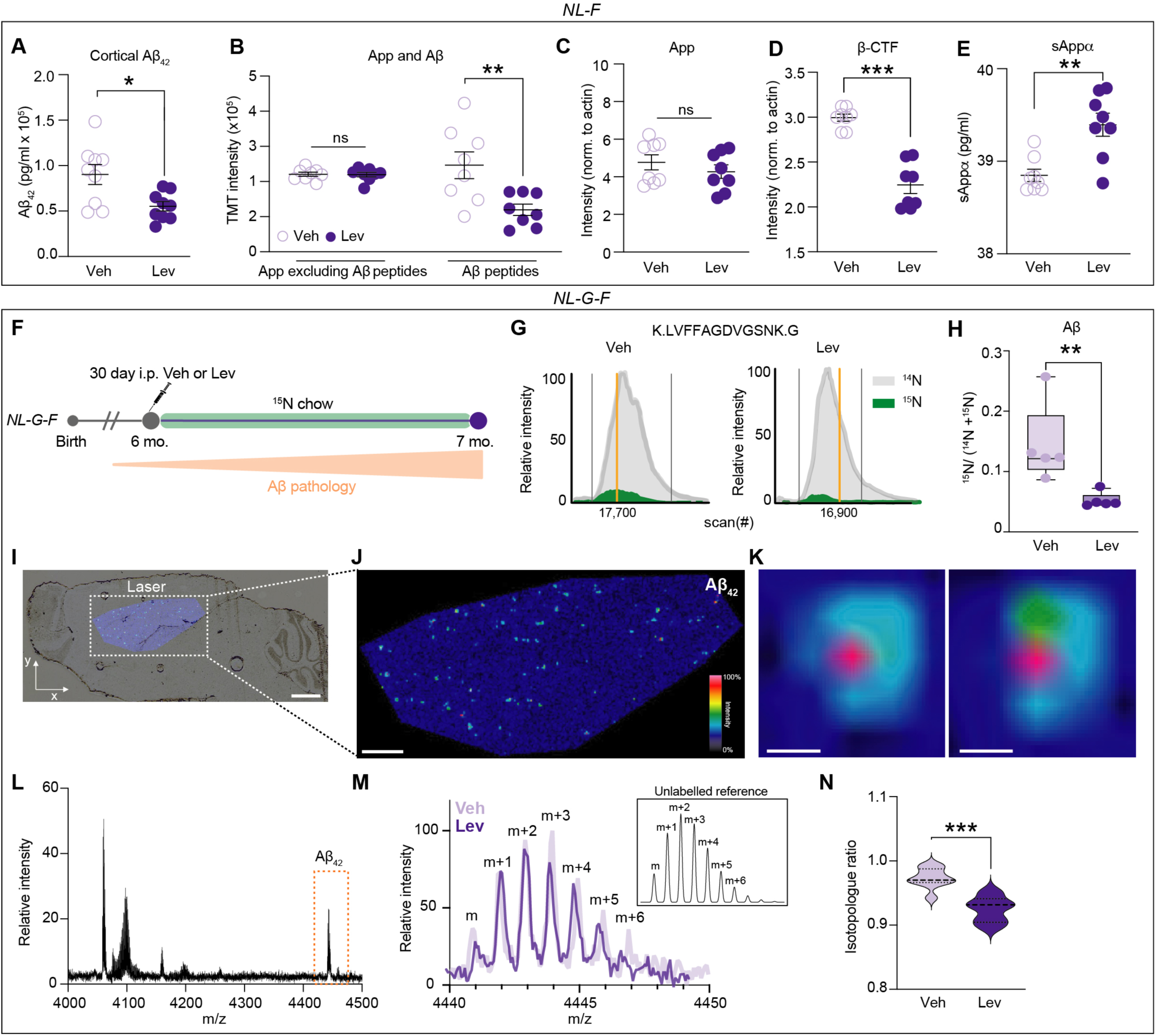
Levetiracetam prevents Aβ_42_ production *in vivo*. (A) ELISA analysis of Aβ_42_ in GuHCl soluble cortical extracts from female *NL-F* mice shows that Lev significantly lowers Aβ_42_ levels. (B) TMT intensities of App peptides mapping outside or within the Aβ_42_ sequence shows that Lev does not alter App levels but does significantly lower the abundance of Aβ peptides in *NL-F* mice. (C-D) Quantification of App WB analysis of *NL-F* cortical extracts confirms that Lev does not affect full length App levels but does significantly lowers β-CTF levels compared to Veh. WBs are presented in Figure S6G and are normalized to actin. (E) ELISA analysis of sAppα, a byproduct of non-amyloidogenic processing, from *NL-F* cortical extracts shows that Lev significantly increases sAppα levels. (F) Schematic depicting metabolic labeling paradigm with ^15^N chow and Lev administration in *NL*-*G-F* mice. Aβ synthesized during Lev treatment will be ^15^N labeled, while Aβ produced before Lev will be fully ^14^N. (G) Representative reconstructed MS1 chromatograms of the Aβ peptide (K.LVFFAGDVGSNK.G) from Lev and Veh cortical 1% SDS insoluble fraction. Grey and green traces indicate relative intensities of the ^14^N and ^15^N ions, respectively. Yellow bar indicates MS/MS scan used to identify the fully ^14^N Aβ peptide with 627.3299 m/z. Back bars indicate the window used for quantification. (H) Quantification of fully ^15^N Aβ relative to total quantified Aβ (^14^N Aβ + ^15^N Aβ) based on the respective reconstructed chromatograms reveals that Lev significantly prevents synthesis of Aβ. (I-J) Matrix-assisted laser desorption/ionization mass spectrometry (MALDI-MS) imaging for Aβ_42_ peptide (m/z 4440.3) across brain sections. Brightfield overlay with representative ion image of Aβ_42_ in section. Intensity scale is normalized ion intensity of Aβ_42_ across region (0-100%). Scale bar for (I) is 1mm and for (J) is 500 µm. (K) Representative single plaques from (J). Scale bar is 25 µm. (L) Plaque region of interest mass spectrum showing Aβ species. Spectrum corresponding to Aβ_42_ is outlined in orange box. (M) Representative spectrum of Aβ_42_ from Veh and Lev used to quantify isotope enrichment from isotopologue ratio. Inset shows isotope spectrum for unlabeled (^14^N) reference. (N) Quantification of (M). Lev animals had significantly less isotoplogue ratios (i.e. less ^15^N Aβ_42_) compared to Veh. (A-E, H) All data are mean ± SEM with n = 5-8 biological replicates. For (M), violin plots are of all amyloid values (5-10 per animal), n = 3 mice (Lev), n = 2 mice (Veh). * = p value < .05; ** = p value < .01; *** = p value < .001; by Student’s t-test for (A, B, C, D, E, H, N)

Our observation that Lev reduces Aβ_42_ levels could occur due to either enhanced clearance or minimized production. Detecting and delineating newly produced from pre-existing Aβ_42_ pools requires a mouse model with aggressive amyloid pathology, therefore we used the *NL-G-F* model. We previously reported that chronic Lev treatment decreases amyloid pathology in *NL-G-F* mice (*27*). To now test if Lev increases clearance or prevents production, we used metabolic ^15^N stable isotope labeling of *NL-G-F* mice to track newly produced Aβ (i.e., ^15^N-labeled) with quantitative MS analysis (**Fig. 5F)**. We quantified the relative peptide abundance of ^15^N Aβ (i.e. ^15^N /(^15^N+^14^N)) with targeted MS and found significantly less newly produced Aβ with Lev treatment (**Fig. 5G-H**). Next, we performed matrix-associated laser desorption/ionization (MALDI)-based MS imaging to visualize and quantify the abundance of ^15^N Aβ_42_ and ^14^N Aβ_42_ from tissue sections of Veh and Lev treated mice. The chemical specificity of this technology allows the spatial quantification of intact Aβ peptides along with relevant isotope content *in situ* (*50*). Specifically, single ion images are generated by mapping the intensity of the Aβ_42_ ion signal (i.e. relative intensity) over the tissue section (**Fig. 5I-K**). The relative abundance of ^15^N Aβ_42_ and ^14^N Aβ_42_ from Veh and Lev treated animals can then be determined based on the isotopologue ratio (**Fig. 5L-M**). With this, we found that Lev significantly decreased the ^15^N Aβ_42_ to ^14^N Aβ_42_ isotopologue ratio compared to Veh treated animals (**Fig. 5N**). These findings demonstrate that Lev decreases Aβ_42_ levels by preventing Aβ_42_ production *in vivo*.

Bulk proteomic analysis of labeled brain extracts additionally provides an opportunity to probe how Lev modulates turnover dynamics in a model we previously discovered had slowed SV protein turnover (*25*). Protein turnover dynamics are quantified using the ^14^N protein fractional abundance (i.e. ^14^N /(^14^N+^15^N)). Lev did not cause a global shift in protein fractional abundance relative to Veh controls (**Fig. S6H)**. GO:CC analysis of the proteins with rescued turnover in Lev compared to Veh groups revealed significantly enriched terms related to presynapse and SVs, include several proteins such as SV2a, Syn1, and Amph **(Fig. S6I)**. To better quantify the effect of Lev on SV2a, we performed GeLC-MS/MS and found that the amount of ^14^N SV2a was significantly reduced by Lev (**Fig. S6J-K**).

Lastly, our earlier findings demonstrated *G76V-GFP/NL-F* mice displayed GFP* accumulation at presynaptic sites, thus we hypothesized that as Lev decreases Aβ_42_, it should consequently decrease GFP* accumulation (**Fig. 1**). To address this, we chronically administered Lev or Veh in *G76V-GFP/NL-F* mice from five to six months and performed subsequent IF analyses. This revealed that Lev significantly reduced GFP* intensity at presynaptic sites compared to Veh controls (**Fig. S6L-N**). Taken all together, these findings confirm that Lev decreases amyloidogenic processing which prevents Aβ_42_ production *in vivo*.

### Levetiracetam prevents synapse deterioration in a transgenic amyloid mouse model

We next addressed whether Lev could minimize synaptic defects *in vivo*. The transgenic *PDGFB-APP^Swe/Ind^ (J20)* mouse model of amyloid pathology was used because *App* KI mice do not model synapse loss (*25, 51, 52*) (**Fig. 6A**). Lev treatment of *J20* mice significantly reduces amyloid pathology-induced cognitive deficits (*32, 53*). To first probe if our findings on the impact of Lev on SVs in *App* KI mice are recapitulated in *J20* mice, we performed electrophysiological patch-clamp recordings from cortical pyramidal neurons in acute brain slices from Veh or Lev treated cohorts. In recordings of miniature excitatory postsynaptic currents (mEPSCs) we did not observe a difference in the amplitude, rise time, or decay times of events in Lev treated mice compared to Veh groups. However, we did measure a significantly reduced frequency of mEPSC events in Lev compared to Veh cohorts (**Fig. 6B-G**). These results indicate that chronic Lev treatment in mice reduces excitatory synaptic transmission.

**Figure 6.**
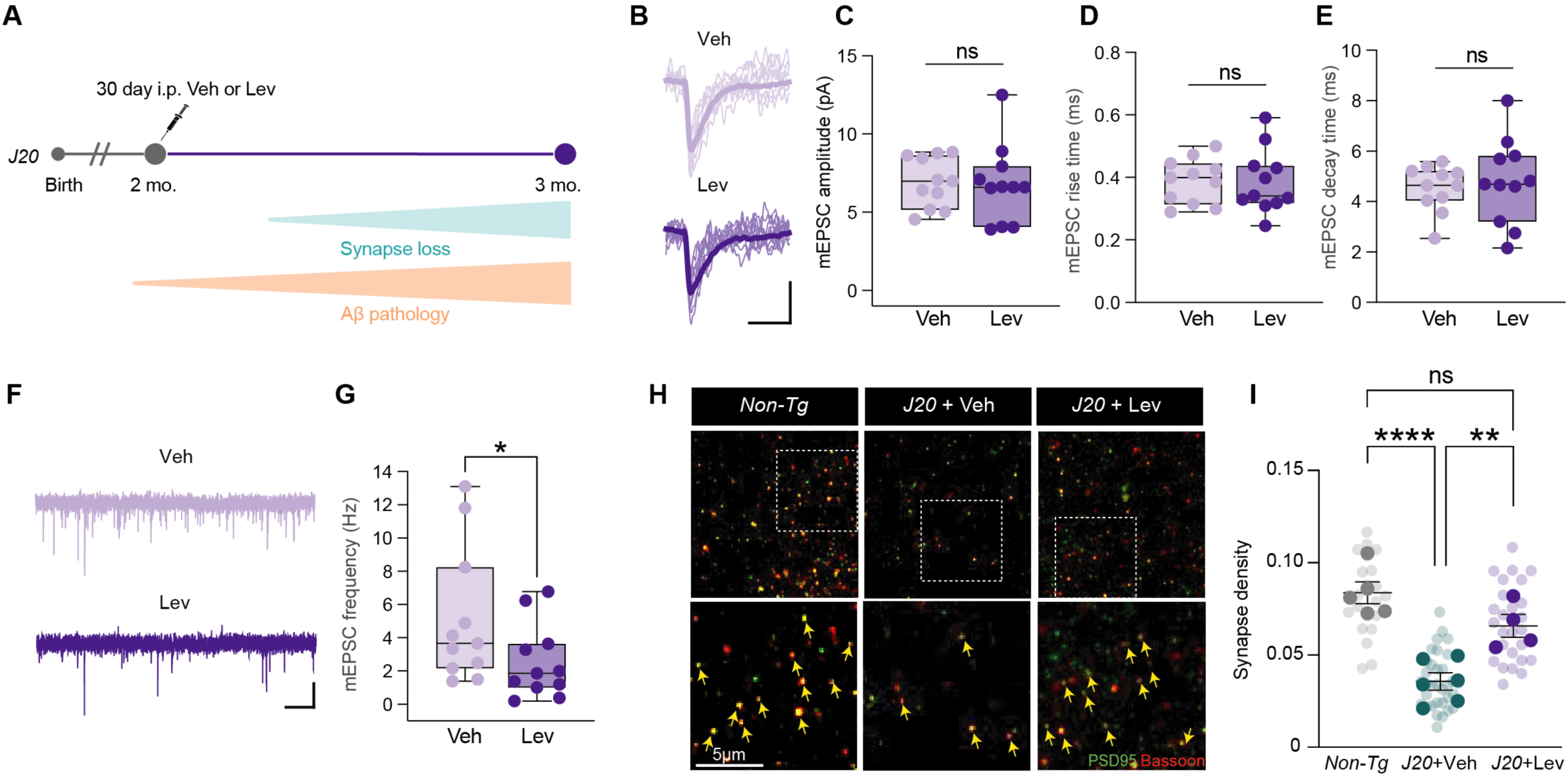
Levetiracetam reduces synapse loss in a transgenic mouse model of amyloid pathology. (A) Schematic depicting pathological timelines and when chronic i.p. administration of Lev or Veh was performed in *J20* mice. (B) Magnified individual mEPSC traces from Veh or Lev treated *J20* mice. 10 traces and average traces are presented in transparent and bold lines. Calibration: 5 ms and 5 pA. (C-E) Collective data of mEPSC amplitude (C), rise time (D), and decay time (E) in individual cells in Veh or Lev treated *J20* mice. (F-G) Collective data of mEPSC frequency in individual cells in Veh and Lev treated *J20* mice. Lev significantly reduced mEPSC frequency compared to Veh. Calibration: 1 s and 10 pA. (H) Representative synapse IF images from Non-Tg, *J20 +* Veh, and *J20* + Lev cohorts. Yellow arrows indicate excitatory synapses. Scale bar is 5 µm. (I) Quantification of (H). Lev significantly rescues synapse density in *J20* mice compared to Veh treatment. Synapse density was defined as number of colocalized Bassoon and PSD95 puncta normalized to area. All data are mean ± SEM with n = 4-6 biological replicates. * = p value < .05; ** = p value < .01; **** = p value < .0001 by one sided Student’s t-test for (C, D, E, and G) or ANOVA with Tukey’s multiple comparisons test for (I).

To verify the timing of synapse loss, we performed IF analysis of brain sections at 1, 2, and 3 months of age in non-transgenic (Non-Tg) and *J20* mice. Consistent with previous findings, no difference in cortical synapse density was detected between Non-Tg and *J20* at 1 or 2 months, but a significant reduction of synapse density in *J20* mice was evident at 3 months (**Fig. S7A-B**) (*52, 54*). Chronic Lev or Veh administration to *J20* mice from 2 to 3 months, followed by quantification of synapse density, revealed Lev significantly minimized synapse loss (**Fig. 6H-I**). These findings show that Lev mitigates synapse loss *in vivo* in an amyloid mouse model.

### Presynaptic proteins accumulate during early stages of Aβ_42_ pathology in human Down syndrome brains

Finally, to evaluate the relevance of Lev treatment to human AD pathology, we sought to determine if human brains highly predisposed for amyloid pathology exhibit elevated levels of presynaptic proteins. Studying the pre-amyloid stages of sporadic AD in humans is challenging because we lack robust AD diagnostic tools needed to conclusively determine which individuals will eventually develop AD (*55*). To overcome this, we studied human Down syndrome (DS) brains, where patients harbor a trisomy of chromosome 21 containing the *APP* gene and have an estimated >90% likelihood of developing amyloid pathology and dementia (*56, 57*). We acquired postmortem DS and control (CTRL) brains from individuals who died at 20-40 years of age (**Fig. 7A**). This represents an important age prior to significant Aβ or amyloid accumulation (*58, 59*). Aβ_42_ and Aβ_40_ ELISA analysis of frontal cortex (FC), entorhinal cortex (EC), and hippocampus (HIP) extracts revealed that Aβ_42_ levels were the highest, although not significant, in the: FC, then EC, and finally in the HIP compared to CTRLs (**Fig. 7B**). This was consistent with previous studies showing that Aβ_42_ pathology begins in the FC before spreading to the EC and HIP in DS patients (*58–60*).

**Figure 7.**
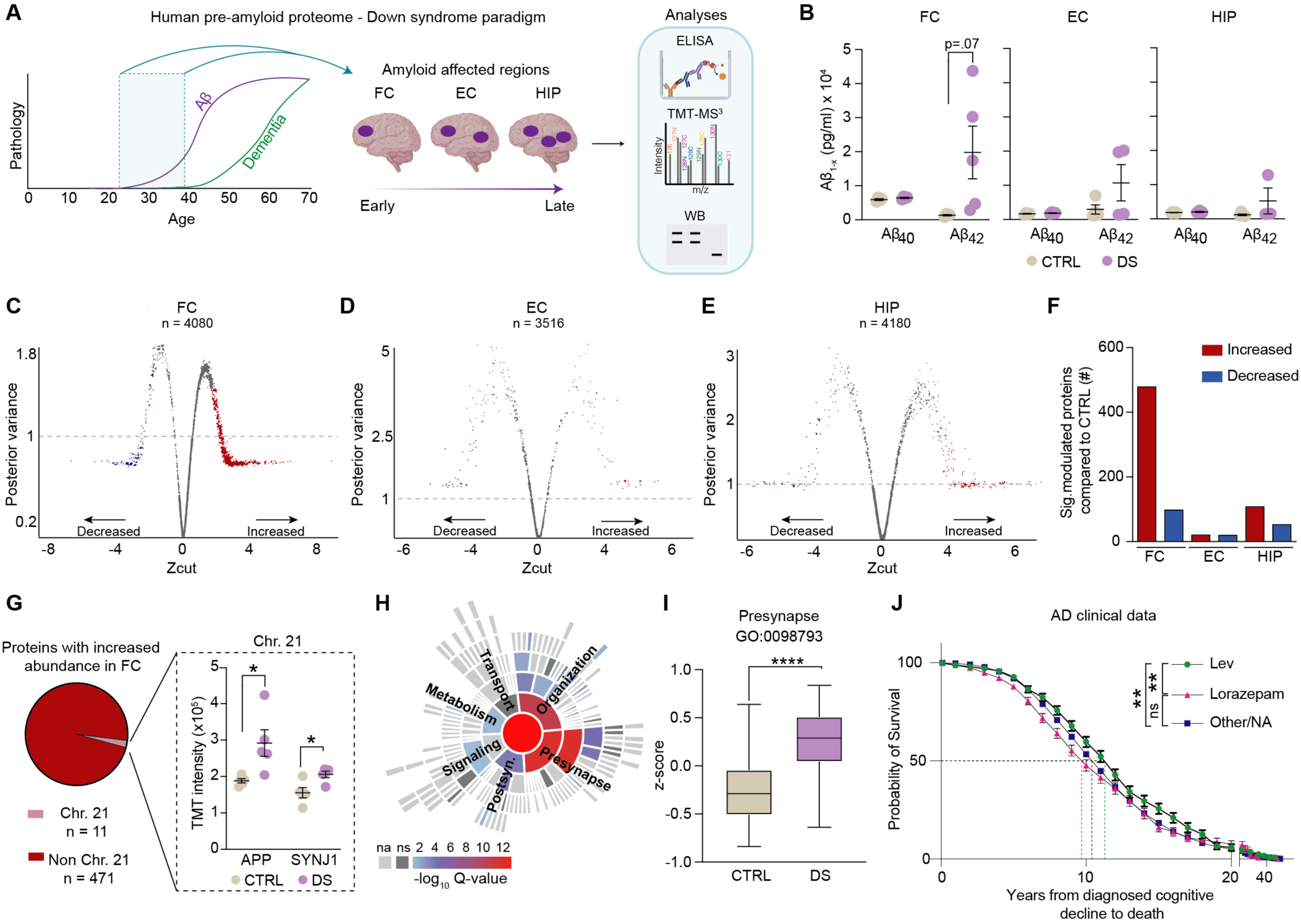
Human Down syndrome brains display presynaptic protein accumulation before significant Aβ_42_ pathology. (A) Schematic depicting the experimental paradigm using human post-mortem Down syndrome (DS) brains. DS and age-matched control (CTRL) brain tissue samples from the frontal cortex (FC), entorhinal cortex (EC), and hippocampus (HIP) between 25-40 years of age (blue box) were analyzed with proteomic and biochemical analyses. (B) Aβ_40_ and Aβ_42_ levels in FC, EC, and HIP GuHCl soluble extracts from DS and CTRL patient samples measured by sandwich ELISA reveal that the FC displays early stages of Aβ_42_ accumulation. (C-E) Shrinkage plots from Bayesian analysis of variance showing protein fold change determined by three TMT-based proteomic experiments of FC, EC, and HIP GuHCl soluble extracts of DS and CTRL samples. The proteins with significantly increased and decreased fold change are in red or blue respectively. (F) Number of significantly modulated proteins in the FC, EC, and HIP from the TMT datasets. (G) Pie chart depicting number of the FC proteins with significantly increased abundance that are encoded on Chr. 21. Inset of TMT intensities from two example Chr. 21 proteins (APP and SYNJ1). (H) SynGO CC analysis of proteins with significantly increased abundance in the DS cohort from the FC dataset showing overrepresentation of the term “Presynapse”. (I) Average TMT intensities of all quantified proteins classified at “Presynapse” with SynGO shows DS FC extracts have significantly elevated levels of presynaptic proteins compared to CTRL. (J) Kaplan-Meier plot showing length of time from diagnosed cognitive decline to death in patients with AD given Lev (n = 472), Lorazepam (n = 634), or other/no antiepileptic drug (n = 26,842). In AD patients, Lev extends time from diagnosis of cognitive decline to death, compared to Lorazepam or no antiepileptics. This analysis utilized data from the Clinical data from the National Alzheimer’s Coordinating Center (NACC). ** = p value < .01 by Mantel-Cox test. All data are mean ± SEM with n=3-8. * = p value < .05; **** = p value < .0001 by Student’s t-test for (B, G, I).

To determine how the brain proteome is remodeled during the pioneering stage of Aβ_42_ pathology, we performed TMT-MS quantitative proteomic analyses (**Fig. S8A-C**). The FC proteome was the most affected compared to the EC and HIP (**Fig. 7C-E, Fig. S8D**). Furthermore, four-fold more proteins had elevated rather than reduced levels, suggesting that Aβ_42_ leads to protein accumulation in human brains (**Fig. 7F)**. We next confirmed that the proteins with elevated levels were not due to increased gene copy number from trisomy 21 and found that in the FC, only 11 of the 482 significantly elevated proteins are encoded on Chr. 21 (**Fig. 7G**). Next, we performed GO:CC enrichment analysis on the significantly elevated proteins, which revealed overrepresented terms such as “axon” and “neuron projection”, with more than a quarter of the proteins associated with synapses (**Fig. S8E**). We next subjected this protein pool to SynGO analysis and found that the GO:CC term “presynapse” was the most significantly overrepresented term (**Fig. 7H-I**). Notably, many proteins involved in SV exo/endocytosis had significantly higher levels in DS compared to CTRL (**Fig S8F**). The phenomenon of elevated levels of SV protein abundance positively correlated with the Aβ_42_ load in DS patient brains (**Fig. S8G**). WB analysis confirmed increased abundance of a panel of these presynaptic proteins in DS brains (**Fig. S8H**). Taken together, these results indicate that human brains highly predisposed for amyloid pathology display presynaptic protein accumulation during the early stages of Aβ_42_ pathology, similar to our previous discovery in *App* KI mice (*25*). Finally, leveraging its status as an FDA-approved and widely used drug, we mined existing human clinical data to investigate whether AD patients who took Lev experienced slowed cognitive decline. To do this, we obtained clinical data from the National Alzheimer’s Coordinating Center and conducted a correlative analysis. Our results, although descriptive, indicate that AD patients who took Lev had a significant delay from the diagnosis of cognitive decline to death compared to those taking lorazepam or no/other AEDs (**Fig. 7J**). While the magnitude of change is small being on the scale of a few years, this analysis supports the positive effect of Lev treatment to slow the progression of AD pathology.

## DISCUSSION

Our findings reveal that presynaptic alterations may represent an important opportunity for therapeutic intervention in AD. Building on our discovery of hampered presynaptic protein degradation before amyloid pathology, we investigated where proteins with impaired degradation build up and found a preferential accumulation at presynaptic sites (*25, 27*). Our biochemical characterization of SVs revealed that Aβ_42_ and the amino terminus of App are in the SV lumen, highlighting the importance of SVs in the establishment of amyloid pathology. The therapeutic potential of targeting SVs to minimize hyperexcitability or reduce amyloid pathology has been demonstrated with small molecule drug Lev (*27, 29, 40*). However, the molecular mechanism by which Lev mitigates amyloid pathology has until now remained elusive. In this study, we discovered that Lev reduces amyloidogenic APP processing by decreasing SV cycling which results in increased surface APP levels. Thus, APP has increased probability to be cleaved by α-secretase, via the non-amyloidogenic pathway. Furthermore, this remarkable effect requires SV2a expression. Finally, we determined that Lev prevents Aβ_42_ production and minimizes synapse loss *in vivo*. These results, in the context of the existing literature, solidifies that targeting SVs represents a promising therapeutic strategy to prevent AD pathology before irreversible damage occurs.

Our study is not without several important limitations. Despite the well documented limitations of using rodents to study AD, these findings highlight that they represent valuable tools to study distinct aspects of AD pathologies (*61*). It is also of note that these models express mutations which cause familial AD and therefore may not fully recapitulate sporadic AD. We additionally acknowledge that tau is an essential aspect of AD pathogenesis and is required for synaptic dysfunction in transgenic APP mice but we did not address this aspect in our study (*62, 63*). This was because the scope of our research was focused on the initial synaptic deficits during the preclinical stage of AD identified in our previous protein turnover study using *App* KI mice. We consistently identified presynapses as the initial site for the manifestation of early Aβ etiology, however tau turnover did not exhibit significant changes at this stage (*25*). The reason for this difference remains unclear and is a key focus of future investigations. Beyond the *App* KI models, our research also utilized DS brains, *J20* mice, and *in vitro* models, all of which overexpress APP. This does lead to the complication that not only are Aβ peptides elevated, but so are all other APP fragments. While DS is often considered a genetic form of AD in which plaques and tangles accumulate and >90% of patients present with dementia, there are some patients who do not develop AD (*57, 64*). Utilizing multiple brain regions from human DS patients with varying Aβ_42_ levels was used delineate proteins likely to accumulate due to Aβ_42_ pathology.

The synaptic deficits during early Aβ accumulation have been unclear, which has hindered the ability to effectively intervene in the pathological trajectory of AD (*65*). We discovered an early impairment in presynaptic protein degradation in AD mouse models and subsequently confirmed that presynaptic proteins accumulate in human DS brains (*25*). This finding is notable as the presynapse is a site where APP proteolytic processing, governed by pH-sensitive secretases, produces Aβ peptides (*8, 10, 14, 16, 17, 49, 66–68*). APP processing is therefore strongly influenced by its localization in membranes or in vesicles (i.e. acidified compartments). Several previous studies have provided evidence that Aβ is physically associated with SVs (*17, 22, 23, 69*). However, our results provide new biochemical evidence from brain extracts that Aβ is present in the lumen of SVs. We and others have also previously found that Aβ peptides can disrupt membrane fusion and SV cycling (*25, 70–72*). Presynaptic perturbations have also been shown to cause β-secretase to accumulate in endosomes, subsequently resulting in increased amyloidogenic processing of APP (*16, 17, 73*). In addition, many genes encoding SV-associated proteins are genetically linked to AD (such as *BIN1* and *PICALM*), further implicating SVs as a substrate of dysfunction (*74–78*). Our findings, taken together with the existing literature, indicate that amyloidogenic processing of APP, in combination with Aβ, results in disrupted SV cycling pathways and excess SV protein accumulation at axon terminals.

Several previous studies have reported that Lev can effectively reduce amyloid pathology and cognitive deficits, and here we have uncovered that this is achieved by restoring non-amyloidogenic processing of APP (*27, 32, 33, 35, 79*). While this finding is novel in the context of AD, Lev has been shown to reduce SV cycling in epilepsy models and when SV2a is overexpressed (*80, 81*). We found that SV2a is required for Lev to prevent Aβ_42_ production and correct SV protein levels. The function of the 12-pass transmembrane protein, SV2a, is not well understood, however recent studies have shown that SV2a recruits and stabilizes Syt1, the principal Ca^2+^ sensor for membrane fusion in the brain (*48, 82*). In our TMT-based proteomic analysis examining the effect of Lev, we found that Syt1 levels are robustly decreased by Lev, an effect that was abolished in the absence of SV2a. Decreased abundance of Syt1 has previously been shown to similarly reduce Aβ levels *in vitro* (*83*). While SV2a is the most reported binding target for Lev, it has also been suggested that Lev reduces synaptic transmission through calcium channels inhibition (*84*). We posit that Lev modulates the SV2a-Syt1 interaction which corrects SV cycling and indirectly results in increased APP levels on the PM.

Our results provide new and compelling evidence that Lev is a strategic therapeutic, capable of preventing the production of Aβ_42_. Notably, because secretases are commonly promiscuous, therapeutics focused on direct secretase modulation have been limited by off-target effects (*85*). Therefore, our discovery that Lev modulates the APP proteolytic processing pathway without directly modulating secretase activity is particularly promising. In terms of new therapeutic opportunities, our results indicate that Lev could be used to delay the onset of amyloid pathology in DS patients. Additionally, Lev could be co-administered with the currently FDA approved amyloid clearing antibodies. Therefore, repurposing Lev to modify AD pathological trajectory offers significant therapeutic opportunity to prevent Aβ_42_ production.

## MATERIALS AND METHODS

### Animals

All experiments performed were approved by the Institutional Animal Care and Use Committee of Northwestern University (IS0009900 and IS00010858 and IS00022178). A total of five mouse models were used: *G76V-GFP* reporter mouse model, three *App* KI mouse models (App^NL/NL^ (*NL*), App^NL−F/NL−F^ (*NL-F*), and App^NL−G−F/NL−G−F^ (*NL-G-F*)) and transgenic *pd PDGFB-APP^Swe/Ind^ (J20)* (*52, 86*). For stable ^15^N isotope labeling, previously described method was followed for *in vivo* labeling (*25, 87–91*). For euthanasia, mice were anesthetized with isoflurane followed by cervical dislocation and acute decapitation. Required brain regions for each experiment were harvested, flash-frozen in a dry ice/ethanol bath, and stored at − 80 °C.

### Human Down syndrome brains

Frozen post-mortem tissues from the frontal cortex, hippocampus, and entorhinal cortex was obtained from UCLA, University of Maryland, University of Pittsburgh, Mt. Sinai, and University of Miami brain banks. Brain tissues were donated with consent from family members of the AD patients and all institutional guidelines were followed during the collection of tissues. Additional details on DS and CTRL patients, their diagnosis, and neuropathological conditions are provided in Table S2.

### SV isolation

Cortical homogenates were diluted with homogenization buffer and centrifuged at 1,000 × g for 15 minutes and the supernatant was collected. The collected supernatant was subsequently spun at 10,000 × g for 15 minutes, and the supernatant was discarded. The pellet (P2) was resuspended in 400 μl of homogenization buffer and the spin was repeated at 10,000 × g for 15 minutes, once again discarding the supernatant. The remaining pellet was resuspended in the 500µl water for hypoosmotic lysis and a glass dounce homogenizer was used to release intact synaptic vesicles. 2µl of 1M HEPES was added to equilibrate the sample before rotation at 4c for 30 min. IZON fractionation was performed with the IZON qEV 35 column and collected into 10 fractions (*92*). Fraction 5 was used to obtain electron micrographs and for WB, LC-MS/MS, Nanoview immunocapture, and proteolysis experiments (*92*). SVs were isolated then incubated with heat deactivated or active Proteinase K to digest cytoplasmic exposed protein domains for 15 min at 37C. This reaction was then quenched with SDS Laemmlli buffer and boiled for 10 min for WB analysis.

### Chronic Levetiracetam administration *in vivo*

Levetiracetam (United States Pharmacopeial) was dissolved in sterile saline solution (0.9% sodium chloride). Equal numbers of male and female mice were randomly assigned to vehicle or Lev groups and were given intraperitoneal (i.p.) injections of 75 mg/kg between 9 am - 12 pm each day for 30 consecutive days (*27*).

### Statistical analysis

Statistical analyses were performed using GraphPad Prism or Orange Data Mining platforms. p-values < 0.05 were considered statistically significant and correction for multiple testing with 5% FDR was performed for non-MS experiments when needed. Hierarchical clustering was performed in Orange to identify clustering. The number of clusters (k) was selected based on optimal silhouette score and minimum 10 or 20 protein group size. Heatmaps are scaled by row (z-score). For Bayesian analysis of variance, we implemented BAMarray 2.0, a Java software package that implements the Bayesian ANOVA for microarray (BAM) algorithm (*93*). The BAM approach uses a special type of inferential regularization known as spike-and-slab shrinkage, which provides an optimal balance between total false detections and total false non-detections.

## Acknowledgments

We thank Dr. Ewa Bomba-Warczak, Dr. Yi-Zhi Wang, Julia Choi, Selene, Sosa, and Cecilia Flores for their contributions and insights. We thank Dr. Clarissa Waites, Dr. Yvette Wong, and members of the Savas laboratory for insightful discussions. The imaging in this article was derived from the Northwestern University Center for Advanced Microscopy, which is generously supported by NCI CCSG P30 CA060553 awarded to the Robert H Lurie Comprehensive Cancer Center. The NACC database is funded by NIA/NIH Grant U24 AG072122. NACC data are contributed by the NIA-funded ADRCs: P30 AG062429, P30 AG066468, P30, P30 AG066509, P30 AG066514, P30 AG066530, P30 AG066507, P30 AG066444, P30 AG066518, P30 AG066512, P30 AG066462, P30 AG072979, P30 AG072972, P30 AG072976, P30 AG072975, P30 AG072978, P30 AG072977, P30 AG066519, P30 AG062677, P30 AG079280, P30 AG062422 P30 AG066511, P30 AG072946, P30 AG062715, P30 AG072973, P30 AG066506 P30 AG066508, P30 AG066515, P30 AG072947, P30 AG072931, P30 AG066546, P20 AG068024 P20 AG068053, P20 AG068077, P20 AG068082, P30 AG072958, P30 AG072959.

## Funding

This work was supported by NIH grants R01 AG078796, R21 AG080248, S10OD032464, and R21 AG080705 and the Cure Alzheimer’s Fund to JNS; NIH grants F31 AG079653 and T32 AG20506 to NRR.; NIH grant RF1 AG022560-16 to RV.

## Author contributions

Conceptualization: NRR and JNS

Methodology: NRR, EFF, RV, AC, and JNS

Investigation: NRR, OD, TN, SL, TJH, JCD, EXD, MD, JG, AU

Writing – original draft: NRR and JNS

Writing – review & editing: NRR and JNS

## Competing interests

The authors declare no competing interests.

## Data and materials availability

The mass spectrometry proteomics data have been deposited to the MassIVE repository with the identifier: (MSV000096225). Further information and requests for resources and reagents should be directed to and will be fulfilled by the Lead Contact, Jeffrey N Savas.

## Notes

### Competing Interest Statement

The authors have declared no competing interest.

