## Supplementary Information and Figures for "Levetiracetam prevents Aβ42 production through SV2a-dependent modulation of App processing in Alzheimer’s disease models"

**SUPPLEMENTARY MATERIALS**

Materials and Methods

Figure S1. Proteins with impaired degradation accumulate at $A\beta_{42}$ and VGluT1 positive synaptic sites.

Figure S2. $A\beta_{42}$ is associated with VGluT1-positive SVs.

Figure S3. Characterization of rat neurons over expressing APP or $APP^{Swe/Ind}$.

Figure S4. Lev lowers β-CTF levels, but not full-length APP, in $APP^{Swe/Ind}$ neurons.

Figure S5. TMT labeling efficiency and enrichment analysis.

Figure S6. Lev corrects elevated levels of presynaptic proteins.

Figure S7. *J20* mice have no change in synapse density at 1 and 2 months.

Figure S8. TMT-MS labeling efficiency and biochemical validation of presynaptic protein accumulation in DS and CTRL brains.

Table S1. Dataset of TMT-MS experiment related to Figure 4 comparing neurons expressing $APP^{Swe/Ind}$ that were Veh, Lev, or SV2a KD + Lev. Relative abundances for all proteins across groups is shown. Significant proteins are shown in Sheet 2.

Table S2. DS and CTRL patient clinical information related to Figure 7.

Table S3. Dataset of TMT-MS experiment related to Figure 7C comparing DS and CTRL FC. Significant proteins are shown in Sheet 2.

### Materials and Methods

#### ELISA assays

$A\beta_{40}$, $A\beta_{42}$, and sAPPβ ELISA analyses were performed in 96-well plates according to the manufacturer's instructions and as previously described (*1, 2*). Human DS brain extracts, *App* KI cortical extracts, and cell lysate were solubilized with 5M GuHCl for 2 h with sonication and vortexing at RT. Solubilized samples were then analyzed with sandwich ELISA according to the manufacturing protocols.

#### Immunohistochemistry experiments

Perfusion, sectioning, and immunohistochemistry procedures were carried out according to previously established methods (*2, 3*). Image acquisition was performed using a Nikon AXR confocal microscope at 63x or CSU-W1-SoRa microscope for super resolution microscopy. Preset thresholding parameters and JACOP plugins were applied for thresholding similarly across experiments and for identification of puncta and to quantify Mander's colocalization coefficients. For synapse density analysis, Bassoon and PSD95 positive puncta were used as a mask and overlaid onto the GFP* channel. When needed, measurement of DAPI or total area was used for analyses to ensure consistent quantification and normalization.

#### Presynaptic and postsynaptic purification

PSD purification was performed using a previously described biochemical method (*4*). In brief, brain tissue was homogenized in Buffer H1 (0.32 M sucrose, in 1 mM $NaHCO_3$, 1mM $MgCl_2$, 0.5 mM $CaCl_2$). Next, the homogenized solution was centrifuged for 20 minutes at 1400 x g. The pellet was washed and dissolved in same buffer H1 and centrifuged at 800 x g for 10 minutes. The supernatant was then centrifuged at 14000 x g for 10 minutes. The pellet was washed and solubilized in Buffer H2 (0.32 M sucrose, 1 mM $NaHCO_3$). Sucrose density gradients containing 0.85, 1.0, and 1.2 M sucrose were prepared and the solubilized pellet was applied on to the top of it. Following ultracentrifugation at 85000 x g for 2 hours, the intermediate layer between 1.2 and 1.0 M sucrose solutions was collected. The synaptosomes-rich fraction was diluted and solubilized mechanically in H2 buffer containing 0.5 % Triton-X and Tris-HCl pH 8.0 and centrifuged at 32500 x g for 30 minutes. The pellet was resuspended in H2 buffer and layered onto the top of already prepared sucrose density gradient with 1.0, 1.5, and 2.0 M sucrose layers. Lastly, ultracentrifugation at 200000 x g for 2 hours enriched PSD membrane components between 1.5 and 2.0 M sucrose layers.

**SV immunocapture**

SVs were analyzed using a custom ExoView Tetraspanin Kit (NanoView Biosciences, USA) with SV detection antibodies, SV2a and Vamp2 along with respective IgGs (*5*). For SV detection, samples were diluted at 1:10,000 then incubated on chips overnight at 4c. Chips were washed three times in solution A and then incubated with a cocktail containing anti-82E1-488, anti-VGAT-568, and anti-VGluT1-647. After 1hr incubation, chips were washed in kit-supplied buffers and imaged by the ExoView R100 using nScan v2.9.5. Data was analyzed using NanoViewer 2.9.5. Fluorescent and colocalization cut-offs were set relative to the IgG control.

**Western blot**

For Western blot analysis, protein concentrations in each sample were determined using the BCA Protein Assay Kit (Thermo Scientific, Cat# 23225). Equal amounts of protein samples were subjected to a five-minute boiling step in SDS Laemmli buffer. Subsequently, the samples were promptly loaded onto either a 10% or 16% gel and subjected to electrophoresis for precise separation of proteins based on their sizes. Proteins were transferred onto a 0.45-micron-sized nitrocellulose membrane. Following a 60-minute blocking period at room temperature (RT), the membranes were incubated overnight at 4 °C with primary antibodies prepared in Tris-buffered saline with 0.1% Tween20 (TBST). The next day, after TBST washing, the membranes were probed with HRP-conjugated secondary antibodies sourced from the same host. Chemiluminescence signals were captured using the Bio-Rad ChemiDoc MP or LICOR Imaging systems.

**Primary neuron culture and lentiviral overexpression of APP and APP$^{Swe/Ind}$**

Primary hippocampal neurons were cultured as previously described (*3*). For lentiviral overexpression of APP and APP$^{Swe/Ind}$ constructs, we used the Syn1-promoter lentiviral backbone deriving from Addgene #30145 and #30137 constructs that had the GFP removed (*6*). At DIV2, the respective lentiviruses were added into the media of primary cultures and left for two days before a full media change. For downstream b and g inhibition experiments, C3 or DAPT was added at DIV14 for 24 hours before RIPA collection and biochemical analyses. For Lev treatment *in vitro*, 150 µM Lev (Abcam) was dissolved in saline solution and added to the media for 24 hours then media. Cell lysate was collected for downstream analyses. For stable isotope labeling with amino acids (SILAC), heavy lysine and heavy arginine (Cambridge Isotopes) were added to media lacking these amino acids. For siRNA knock down of SV2a, SV2b, and non-targeting controls, 6-8µl of respective siRNA from (Horizon Discovery) was added to the media of lentivirus infected neurons at DIV11 and left for 4 days for efficient knock down.

**Biotin labeling and Neutravidin pull-down of surface proteins**

Primary neurons treated with either Lev or Veh for 24 hours were moved to a cold room (4 °C) and washed twice with ice-cold PBS. Primary amines at the cell surface were labeled with sulfo-NHS-SS-biotin (Sigma-Aldrich) in ice-cold PBS for 30 min under gentle shaking. The non-reacted sulfo-NHS-SS-biotin was quenched for 2 × 3 min with a buffer containing 65 mM Tris, 150 mM NaCl, 1 mM CaCl2, and 1 mM MgCl2 at pH 7.5, and the cells were washed twice more with ice-cold PBS. Neurons were then collected in RIPA buffer with the addition of protease inhibitors. Biotinylated proteins in the sample were then purified using neutravidin beads. The recovered biotinylated proteins were eluted with SDS Laemmli buffer and prepared for WB analysis.

**Surface APP ICC**

Primary neurons on coverslips treated with either Lev or Veh for 24h were moved to 4°C for 5 min then an 4 µl of N-terminal APP antibody was added to the media for 15 min. Neurons were washed (3 × 5 min) with ice cold PBS before fixation with 4% PFA for 15 min at RT. Fixed neurons were then washed (3 × 5 min) with PBS then blocked in 10% HS with no detergents for 3h. Primary antibody for TransferrinR were then added in PBS overnight at 4°C. The next day, sections were washed with PBS (3 × 5 min) and then incubated with secondary antibodies in

PBS. After secondary antibody incubation, sections were washed with PBS (3 × 5 min) and coverslips were mounted with Fluoromount-G. Images were taken using a Nikon AXR confocal microscope at 63x.

**Syt1-antibody assay**

Primary neurons on coverslips treated with Lev or Veh for 24h were utilized for this assay. 4 µl Syt1luminal-647 (Synaptic Systems 105 311AT647N) antibody was added directly to the media then kept at 37°C for 3 min. Neurons were washed (3 × 5 min) with media before fixation with 4% PFA for 15 min at RT in the dark. Fixed neurons were then washed with PBS then blocked in 0.2% Triton-X 100 and 10% Horse Serum (HS) in PBS for 3h at RT. Primary antibodies against Syt1luminal and Syp were then added in diluted blocking buffer overnight at 4°C. The next day, coverslips were washed with PBS (3 × 5 min) and then incubated with secondary antibodies in PBS. After secondary antibody incubation, sections were washed with PBS (3 × 5 min) and coverslips were mounted with Fluoromount-G. Images were taken using a Nikon AXR confocal microscope at 63x.

**Electrophysiology**

Single-cell patch clamp recording were made from acutely prepared brain slices  from *J20* mice that had been chronically treated with Veh or Lev (75 mg/kg i.p.). Mice were sacrificed then used for experiments same day. The brain was removed, blocked and coronal brain slices (300 µm thick) were made using a vibrating blade microtome (Leica VT1200) while keeping the tissue equilibrated in ice-cold sucrose based artificial cerebrospinal fluid (ACSF) containing (in mM): 85 NaCl, 2.5 KCl, 1.25 NaH2PO4, 25 NaHCO3, 25 glucose, 75 sucrose, 0.5 CaCl2, and 4 MgCl2 with 10 µM DL-APV and 100 µM kynurenate. Slices were incubated for ~15 min in the same sucrose based ACSF at 30 °C, then were allowed to recover at room temperature for at least 1.5 h while the solution was gradually exchanged for a recovery ACSF containing (in mM): 125 NaCl, 2.4 KCl, 1.2 Na2PO4, 25 NaHCO3, 25 glucose, 1 CaCl2, and 2 MgCl2 with 10 µM

DL-APV and 100 μM kynurenate. During recordings, cells were visualized under Dodt contrast optics and were perfused with normal ACSF containing (in mM): 125 NaCl, 2.4 KCl, 1.2 Na2PO4, 25 NaHCO3, 25 glucose, 2 CaCl2, and 1 MgCl2. Solutions were continuously equilibrated with 95% O2 and 5% CO2 throughout experiments. Voltage-clamp recordings were made from pyramidal neurons located in L5 of the somatosensory cortex using standard visualized patch-clamp techniques. Recording electrodes had tip resistances of 3 - 5 M□ when filled with internal recording solution containing (in mM): 135 CsMeSO3, 10 TEA-Cl, 0.2 EGTA, 10 HEPES, 1 MgCl2, 10 Na-phosphocreatine, 2 Mg-ATP, 0.3 Na-GTP, and 5 QX-314. After forming a GΩ seal the whole-cell configuration was established and neurons were voltage-clamped at -70 mV. Miniature EPSCs (mEPSCs) were isolated by perfusing the slice with ACSF including the GABAA receptor antagonist picrotoxin (PTX, 50 μM) and sodium channel antagonist tetrodotoxin (TTX, 0.5 μM). Data were collected and analyzed using pClamp 10 software (Molecular Devices).

**MALDI-MS Imaging**

Fresh frozen tissue sections on ITO-coated glass slides were fixed in 100% ethanol for 60 seconds, followed by 70% ethanol for 30 seconds. Lipids were removed by immersing the sections in Carnoy's solution (6:3:1 ethanol/chloroform/acetic acid) for 110 seconds, followed by subsequent washes in 100% ethanol for 15 seconds, 0.2% trifluoroacetic acid in water for 60 seconds, and 100% ethanol for 15 seconds. The tissues then underwent peptide retrieval using formic acid vapor for 20 minutes. To facilitate peptide ionization and desorption, a matrix solution (15 mg/ml 2,5-dihydroxyacetophenone in 70% acetonitrile, 2% acetic acid, 2% trifluoroacetic acid) was applied using a TM sprayer (HTX Technologies, NC, USA) and recrystallized in 5% methanol vapor at 85°C for 3 minutes. MALDI-MSI was performed using a MALDI-time-of-flight (TOF) instrument (RapifleX, Bruker Daltonics) equipped with a scanning Smartbeam 3D laser. The acquisition was conducted at a pixel size of 20 μm with 200 shots per pixel at 90% power with a shot frequency of 10 kHz. Spectra were collected within the mass range of 1500–6000 m/z in reflector positive mode (mass resolution: m/Δm=15000 (FWHM) at m/z 4440.3). The system was externally calibrated by spotting a mixture of peptide standard II and protein standard I (Bruker Daltonics). Plaque ROI annotation, and ROI spectra export as *csv was performed using the FlexImaging 5.1 software (Bruker Daltonics). The ROI spectral data were processed ana analyzed in R by means of baseline substraction and RMS normalization. Relative peak areas for the third and fourth isotopologue (4442.3 and 4443.3) were calculated. Isotope content was calculated by comparing the m/z 4442.3/ m/z 4443.3 ratio ($R_m$) of measured A$\beta_{42}$ (arc) vs a theoretical calculated value predicted for the non-labelled species containing only natural abundance isotope content ($R_{std}$).

**TMT-MS sample preparation**

TMT-MS sample preparation was performed as previously described (*1, 3, 7*). Isobaric TMT labeling of 100 µg peptides from each sample was performed according to manufacturer instructions. After a 60-minute incubation at room temperature, the reaction was quenched with 5% hydroxylamine. Isobarically labeled samples were combined in equal proportions, quickly desalted, and fractionated into 8 fractions using high pH reversed-phase columns. The resulting peptide solutions were dried, stored at −80 °C, and reconstituted in LC-MS Buffer A (5% acetonitrile, 0.125% formic acid) for LC-MS/MS analysis.

**TMT data collection**

TMT-MS3 analysis was performed as previously described (*1, 3, 7*). The Orbitrap Fusion was employed for MS data generation using a Multinotch MS3 method. In the MS3 channel, the top ten precursor peptides were selected for SPS-MS3 analysis and fragmented using 65% HCD before orbitrap detection. Precursor selection ranged from 400–1200 m/z with mass range tolerance. Exclusion mass width was set to 18 ppm low and 5 ppm high. Isobaric tag loss exclusion was applied to TMT reagent. Additional MS3 settings include an isolation window = 2, orbitrap resolution = 60 K, scan range = 120–500 m/z, AGC target = 6*105, max injection time = 120 ms, microscans = 1, and datatype = profile.

**LC-MS/MS and TMT-MS sample preparation**

Samples were prepared for LC-MS/MS analysis as previously described (*3, 7*). In short, proteins were precipitated using methanol/chloroform precipitation, denatured with 8 M urea, and subsequently processed with ProteaseMAX. TMT-MS sample preparation was performed as previously described (*1, 3, 7*). Isobaric TMT labeling of 100 µg peptides from each sample was performed according to manufacturer instructions. Isobarically labeled samples were combined in equal proportions, quickly desalted, and fractionated into 8 fractions using high pH reversed-phase columns.

**LC-MS/MS and TMT-MS data analysis**

Protein identification, quantification, and analysis utilized the Integrated Proteomics Pipeline - IP2 (Integrated Proteomics Applications, Inc., San Diego, CA, http://www.integratedproteomics.com/) as previously described, employing ProLuCID, DTASelect2, Census, and QuantCompare (*2, 3, 7*). LC-MS/MS experiments involved extracting spectrum raw files into MS1, MS2 files using RawExtract 1.9.9 (http://fields.scripps.edu/downloads.php). ProLuCID searches included fully and half-tryptic peptides within the mass tolerance window with unlimited miscleavages. Static modification considered carbamidomethylation (+57.02146 Da) of cysteine. DTASelect2 assessed PSMs using XCorr and DeltaCN. Peptide probabilities and FDR were calculated based on a target/decoy database with proteins having an FDR of ≤1% at the protein level and requiring a minimum one peptide of six amino acid residues. For dynamic $^{15}N$ experiments, datasets were searched against light ($^{14}N$) and heavy ($^{15}N$) protein databases separately (*8, 9*). Fractional abundance was calculated from the $^{14}N$ and $^{15}N$ peptide chromatograms, and for each respective protein the $^{14}N$ and $^{15}N$ chromatograms for the respective peptides was averaged. For TMT-MS experiments, data analysis was performed as previously described (*7*). Protein identification required a minimum peptide length of six, with each dataset having a 1% FDR rate at the protein level. Isobaric labeling analysis was conducted with Census2 without applying an intensity threshold.

**Gene ontology enrichment analysis**

For Gene ontology analysis, we used protein analysis through: ShinyGO, Pantherdb, or SynGO to determine overrepresented cellular component categories (*10*). The statistical overrepresentation test was calculated by using the significant proteins identified from respective groups as the query and the aggregated total proteins identified for the respective experiment as the reference. Ontologies statistical tests with FDR correction less than .01 were considered significant.

### Supplementary Figures and Legends

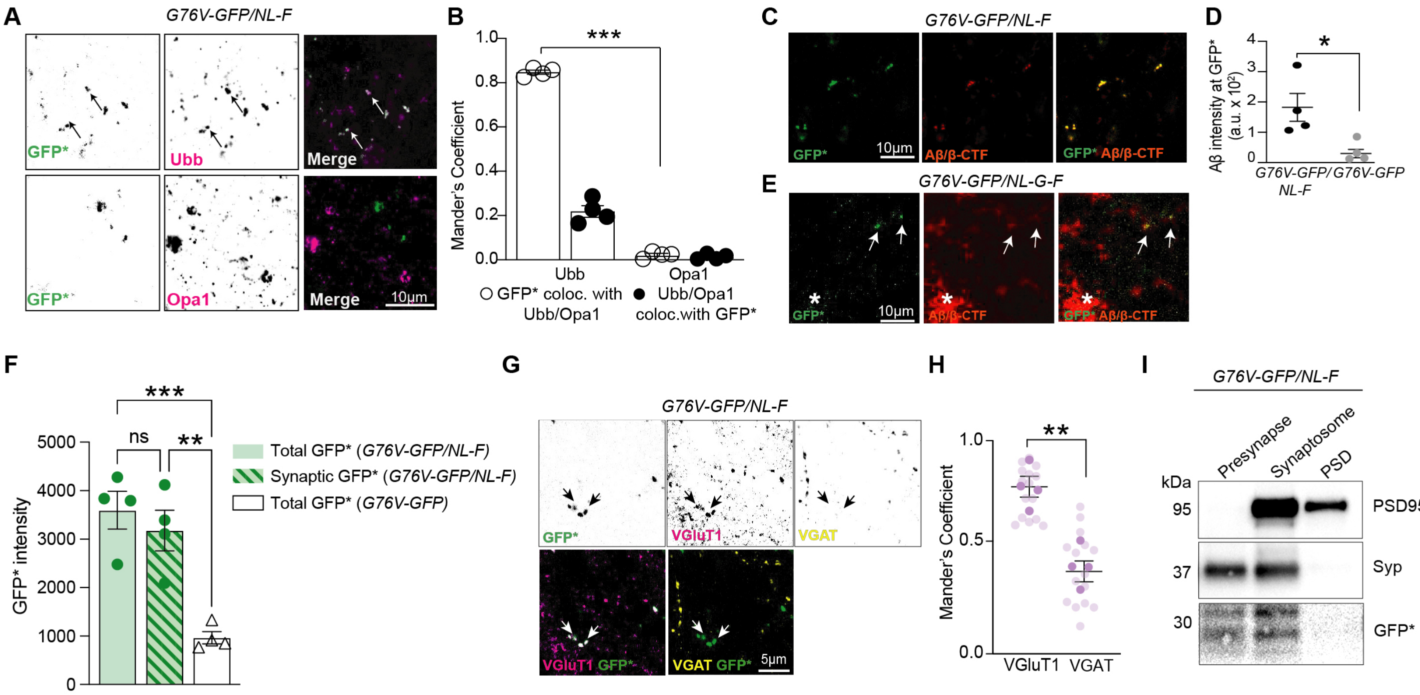


**Figure S1. Proteins with impaired degradation accumulate at $A\beta_{42}$ and VGluT1 positive synaptic sites.**

(A) Representative IF images showing GFP* colocalization with Ubb but not Opa1 puncta in *G76V-GFP/NL-F*. Scale bar is 10µm.

(B) Quantification of (A). Mander's correlation coefficient between GFP* and Ubiquitin puncta (Ubb) is significantly higher than GFP* with mitochondrial protein Opa1 in *G76V-GFP/NL-F*.

(C) Representative IF images showing GFP* colocalization with Aβ (82E1 antibody) puncta in *G76V-GFP/NL-F*. Scale bar is 10µm.

(D) Quantification of (C). *G76V-GFP/NL-F* mice have significantly higher intensity of Aβ (82E1 antibody) at GFP* puncta in the cortex compared to control mice.

(E) Representative IF analysis of 6 month *G76V-GFP/NL-G-F* mice demonstrates GFP* is present at Aβ puncta (arrows) but does not colocalize with large Aβ deposits (star). Scale bar is 10µm.

(F) Quantification of total GFP* and synaptic GFP* intensity from *G76V-GFP/NL-F* and total GFP* intensity from *G76V-GFP* shows that over 90% of the total GFP* intensity is synaptic.

(G) Representative IF images show that GFP* colocalizes with VGlut1 puncta but not VGAT puncta in *G76V-GFP/NL-F*. Scale bar is 5µm.

(H) Quantification of (F). Mander's correlation coefficient of GFP* with excitatory presynaptic puncta (VGluT1) is significantly higher compared to GFP* with inhibitory presynaptic puncta (VGAT) in *G76V-GFP/NL-F*.

(I) Biochemical isolation of synaptosome, presynaptic, and postsynaptic compartments from *G76V-GFP/NL-F* cortical homogenates confirms that GFP* is detected in the synaptosome and presynaptic but not postsynaptic fraction. All data are mean ± SEM with n = 4 mice at 6 months of age. * = p value < .05; ** = p value < .01; *** = p value < .001; by Student's t-test for (B, D H) or ANOVA with Tukey's multiple comparisons test for (F).

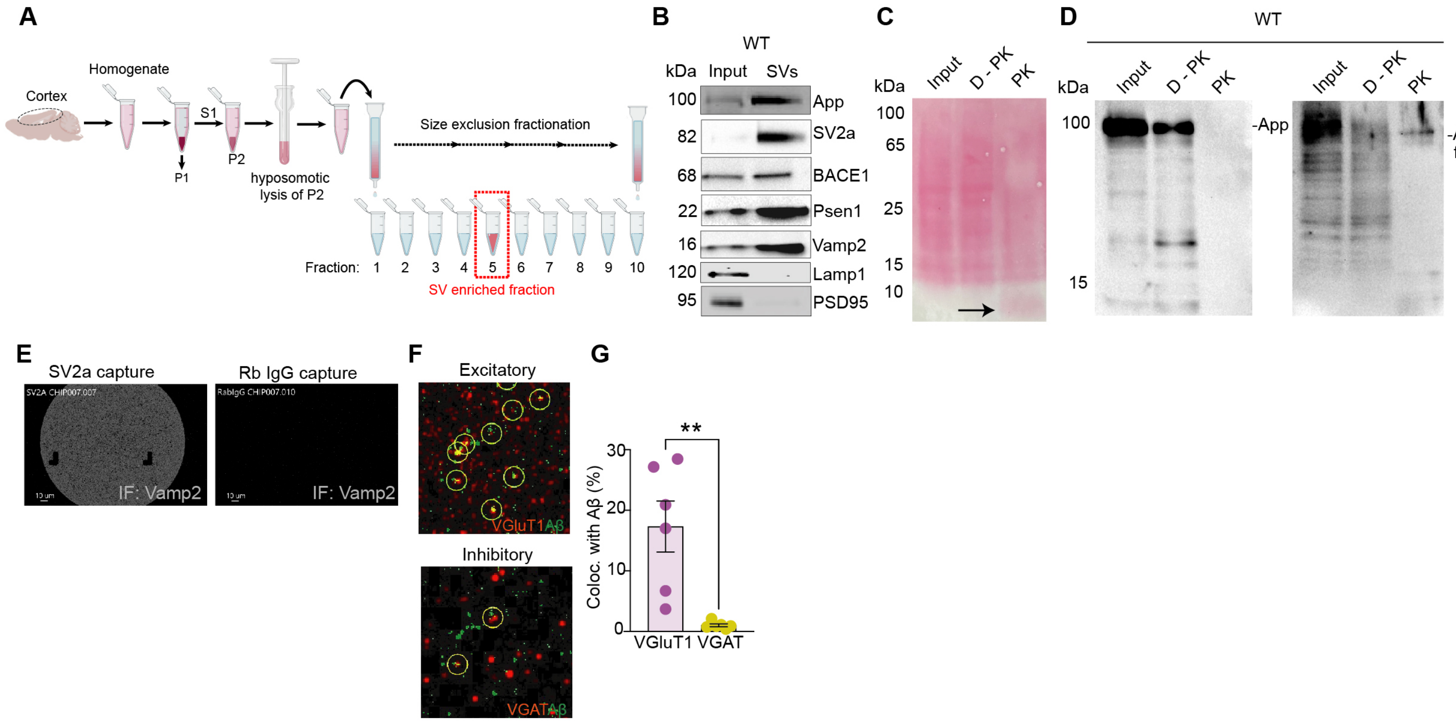


**Figure S2. A$\beta_{42}$ is associated with VGluT1-positive SVs.**

(A) Schematic depicting Proteinase K (PK) treatment of isolated SVs used to probe the orientation of App with Western blot analysis.
(B) Representative WB analysis of F5 showing SV enrichment with hypoosmotic lysis combined with size-exclusion column (IZON) strategy from WT cortical homogenates.
(C) PonceauS staining of total protein load confirms that PK treatment effectively digests proteins (arrow).
(D) WB analysis of SVs (Input), SVs treated with deactivated PK (D-PK), and SVs treated with PK from WT extracts probed with a C-terminal App antibody shows that no C-terminal App is present after PK treatment, indicating its cytoplasmic orientation. N-terminal App antibody shows that N-terminal App is present at (~90 kDa) after PK treatment, indicating its luminal orientation.
(E) Representative intact SV immunocapture with SV2a or Rabbit IgG antibodies followed by immunostaining with Vamp2 (white) show specific capture of intact SVs.
(F-G) SV immunocapture of isolated SV from *NL-G-F* cortical extracts combined with immunostaining with VGluT1, VGAT, and $A\beta_{42}$ showed that $A\beta_{42}$ positive SVs were significantly more colocalized with VGluT1 than VGAT. All data are mean ± SEM with n = 6 mice. ** = p value < .01; by Student's t-test for (G).

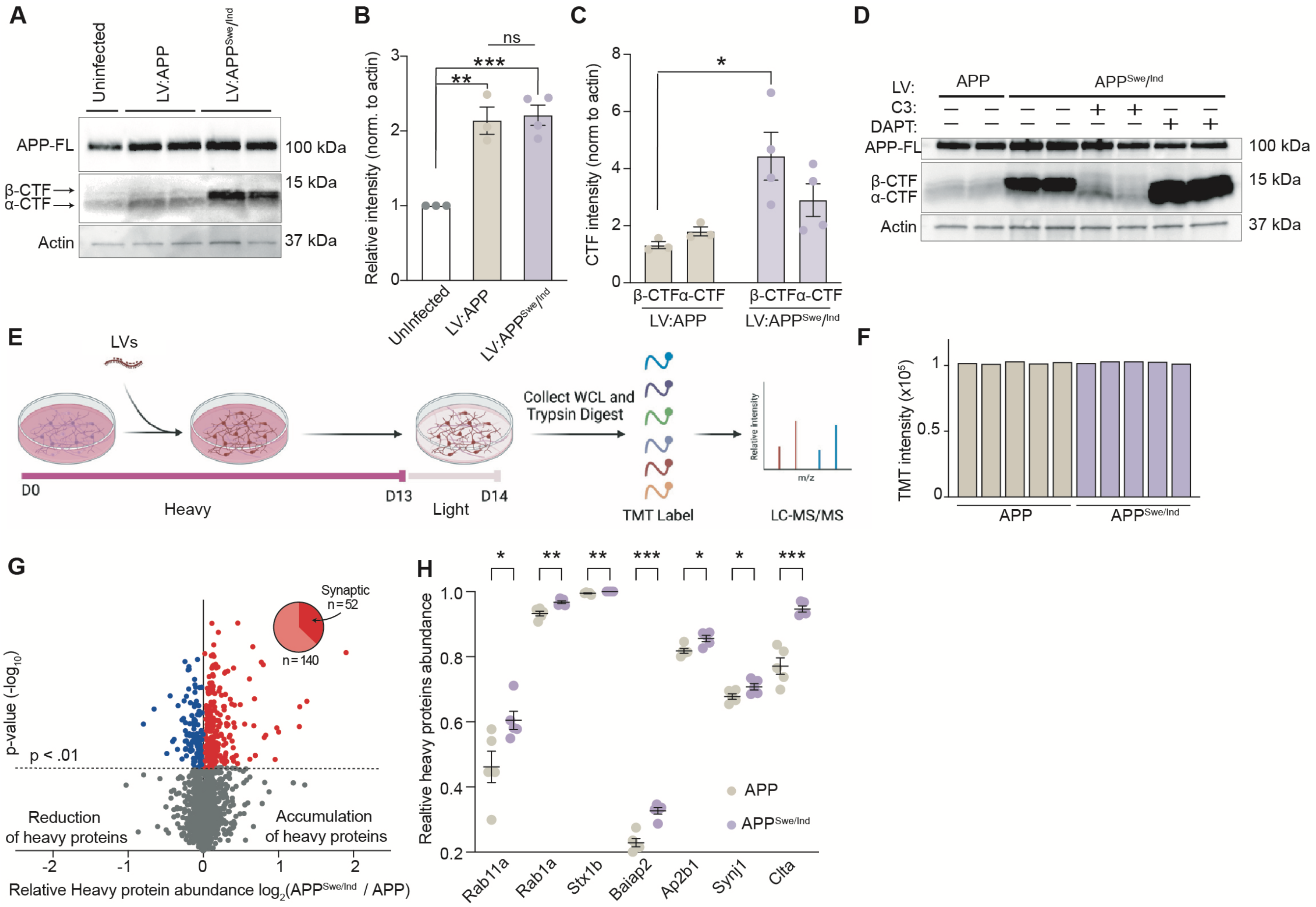


**Figure S3. Characterization of primary neurons overexpressing either APP or APP$^{Swe/Ind}$.**

(A) Representative WB of neurons infected with lentiviruses overexpressing APP or APP$^{Swe/Ind}$.

(B) Quantification of (A). Neurons overexpressing APP and APP$^{Swe/Ind}$ both have a two-fold significantly higher level of full-length APP compared to uninfected neurons. WB analysis normalized to actin.

(C) Quantification of (A). APP$^{Swe/Ind}$ expressing neurons have significantly higher β-CTF levels compared to APP neurons. WB analysis normalized to actin.

(D) Representative WB analysis confirms that APP processing can be modulated with small molecule β– (C3) and γ– (DAPT) secretase inhibitors.

(E) SILAC labeling with TMT-MS analysis schematic to investigate turnover dynamics in APP and APP$^{Swe/Ind}$ neurons. Protein synthesized from DIV1-13 incorporate the Lys+6 and Arg+10 amino acids (Heavy) while proteins synthesized from DIV13-14 incorporate light amino acids (Light).

(F) Global TMT channel peak intensities for each of the 10-plex TMT reporter ions determined from MS analysis demonstrates equal labeling across all channels.

(G) Volcano plot depicting relative heavy protein ($\geq$ one day old) levels in $APP^{Swe/Ind}$ compared to APP expressing neurons. Blue and red features indicate reduced or elevated proteins with p value < 0.05. Pie chart shows proportion of red proteins that are synaptic. N = 3732 proteins.

(H) Selected presynaptic proteins with significantly more relative heavy protein ($\geq$ one day old) levels in $APP^{Swe/Ind}$ compared to APP expressing neurons.

All data are mean ± SEM with n=3-5. * = p value < .05; ** = p value < .01; *** = p value < .001; by Student's t-test for (H) or ANOVA with Dunnet's multiple comparisons test for (B, C).

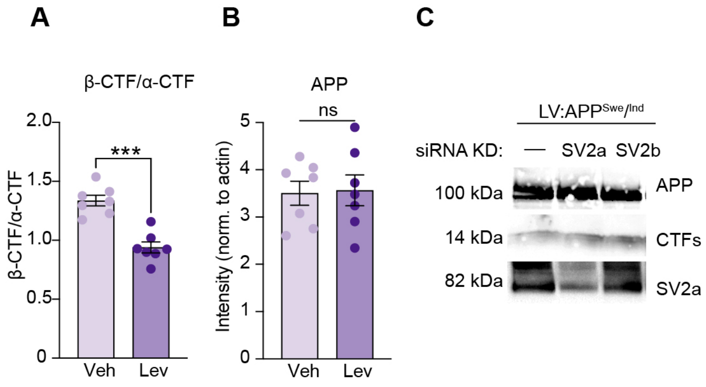


**Figure S4. Lev lowers β-CTF levels, but not full-length APP, in APP$^{Swe/Ind}$ neurons.**

(A) Quantification of WB from Figure 3B shows that the ratio of β-CTF to α-CTF is significantly decreased in APP$^{Swe/Ind}$ neurons treated with Lev compared to Veh.

(B) Quantification of WB from Figure 3B shows no significant differences in abundance of full-length APP in APP$^{Swe/Ind}$ neurons treated with Lev compared to Veh. WB analysis normalized to actin.

(C) Representative WB analysis of APP$^{Swe/Ind}$ neurons with SV2a KD and SV2b KD shows no alteration in APP or β-CTF levels. All data are mean ± SEM with n=7. *** = p value < .001by Student's t-test for (A, B).

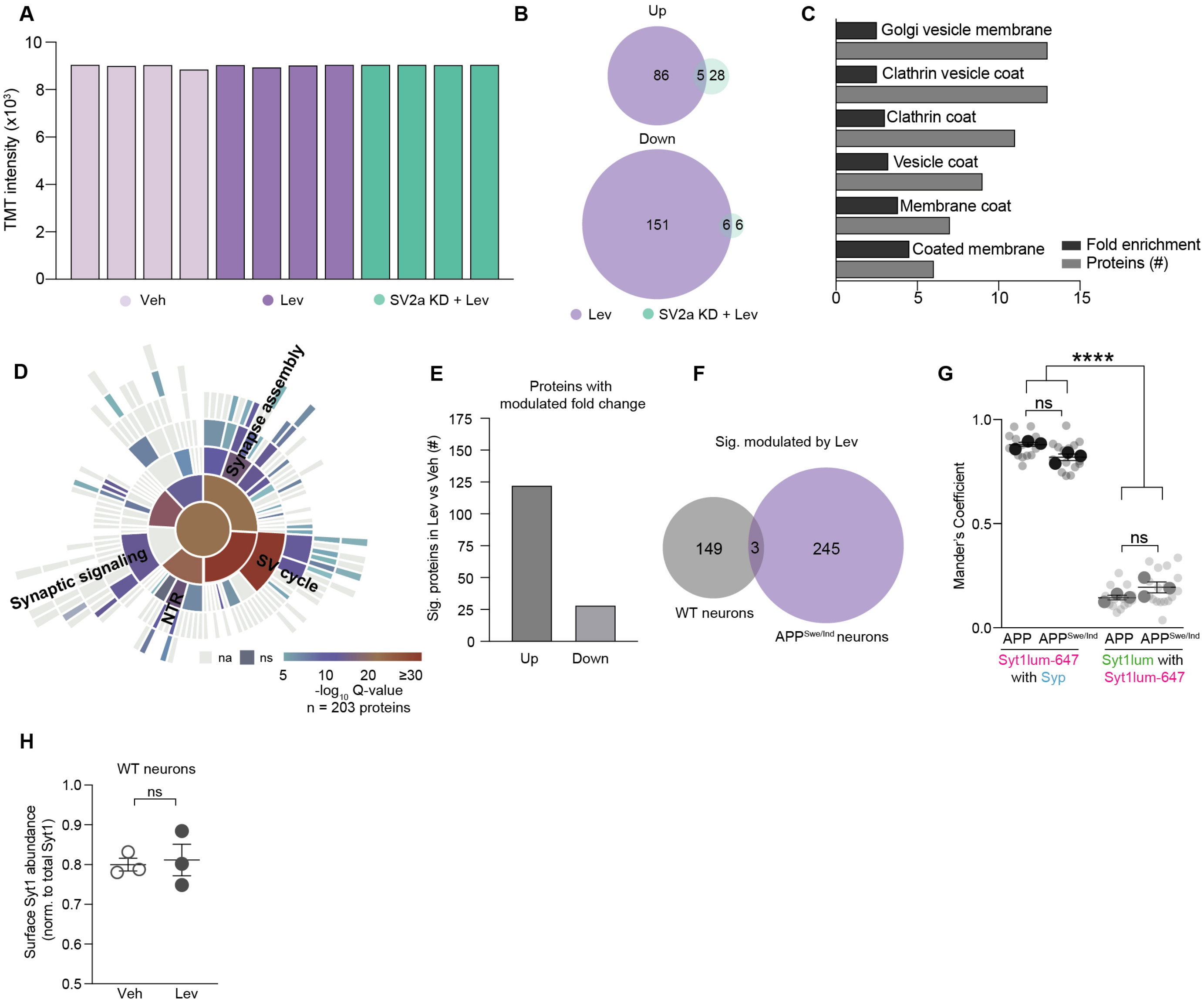


**Figure S5. TMT labeling efficiency and enrichment analysis.**

(A) Representative global TMT channel peak intensities for each reporter ion from 12-plex TMT-MS demonstrates equal labeling across all channels.

(B) Venn diagram showing very few of the significantly modulated proteins from the Lev or SV2a + Lev comparison to Veh (from Fig. 4B) are actually modulated in both datasets.

(C) GO:CC analysis showing the cellular components that are overrepresented among the pool of significantly modulated proteins in the Lev compared to Veh dataset.

(D) Sunburst SynGO plot showing the most significantly enriched cellular component term is "SV Cycle" based on the proteins with decreased abundance relative to both Veh and SV2a KD + Lev (n = 203 proteins)

(E) Summary of proteins with significantly altered fold change from TMT-MS analyses of WT neurons treated with Lev versus Veh with n = 5 for each group.

(F) Venn diagram comparing the proteins with significantly altered fold change in the WT or $APP^{Swe/Ind}$ datasets shows that only three non-synaptic proteins are altered in both. This indicates that Lev does not modulate synaptic proteins in WT neurons.

(G) Mander's correlation coefficient between Syt1lum-647 and Syp intensities in APP and $APP^{Swe/Ind}$ over-expressing neurons (~0.85 - 0.95) was not significantly different. The Mander's coefficient between Syt1lum and Syt1lum-647 in APP and $APP^{Swe/Ind}$ over-expressing neurons (~0.15 - 0.20) highlights the low degree of colocalization and was not significantly different. This indicates the paradigm effectively delineates the two discrete pools of Syt1.

(H) Ratio of Syt1lum-647 surface puncta to total Syt1 puncta (Syt1lum-647 + Syt1lum) from Veh and Lev treated WT neurons. Lev treatment did not alter surface Syt1 (Syt1lum-647) relative to total Syt1. All data are mean ± SEM with n = 3-5 independent biological replicates. **** = p value < .0001 by Student's t-test for (H) or ANOVA with Sidak multiple comparisons test for (G).

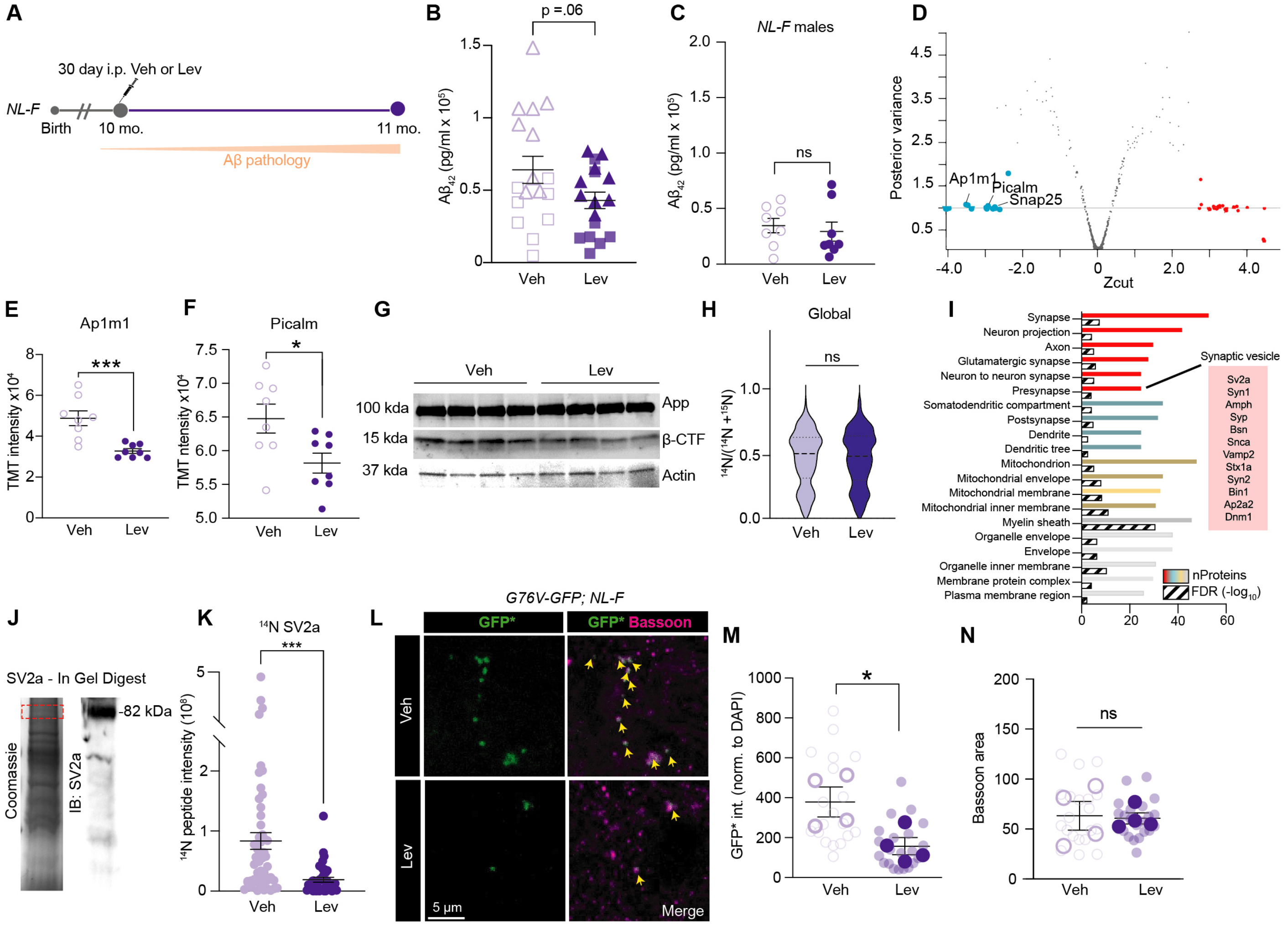


**Figure S6. Lev corrects the elevated levels of presynaptic proteins.**

(A) Schematic depicting timeline of chronic i.p. administration of Lev or Veh to *NL-F* mice.

(B) A$\beta_{42}$ ELISA analysis shows that Lev decreases A$\beta_{42}$ levels in *NL-F* mice compared to Veh. Female and male animals are shown with triangles and squares respectively, highlighting the clear separation of the two sub-populations.

(C) A$\beta_{42}$ ELISA analysis shows a slight non-significant decrease in A$\beta_{42}$ levels in male *NL-F* mice treated with Lev compared to Veh.

(D) Shrinkage plot showing proteome-wide differences of female mice receiving Lev or Veh. Red indicates proteins with significantly elevated levels by Lev. Blue indicate proteins with significantly decreased levels by Lev.

(E-F) TMT peptide intensities from selected presynaptic proteins shows that Lev significantly decreases levels compared to Veh.

(G) Representative App and β/α-CTFs WBs from Lev and Veh treated female *NL-F* mice.

(H) Lev does not affect average global protein fractional abundance (i.e., protein turnover) in *NL-G-F* cohorts compared to Veh.

(I) Proteins with rescued turnover with Lev were subjected to GO:CC analysis. GO:CC terms such as 'Presynapse', and 'Synapse' were among those significantly overrepresented.

(J) Coomassie stained SDS-gel and SV2a immunoblotting used to select and excise the appropriate molecular weight band for GeLC-MS analysis.

(K) GeLC-MS analysis showed that Lev significantly decreases the level of fully $^{14}N$ SV2a peptides.

(L) Representative IF images from *G76V-GFP;NL-F* mice treated with Veh or Lev from 5-6 months stained with Bassoon antibodies.

(M-N) Quantification of (L). Lev significantly reduces the intensity of GFP* at Bassoon positive puncta in *G76V-GFP; NL-F* mice. GFP* intensity was normalized to the DAPI. No significant difference in the total area Bassoon puncta was quantified between Veh and Lev treated *G76V-GFP; NL-F* mice. All data are mean ± SEM with n = 4-8 biological replicates. * = p value < .05; *** = p value < .001; by Student's t-test for (B, C, E, F, H, K, M, N).

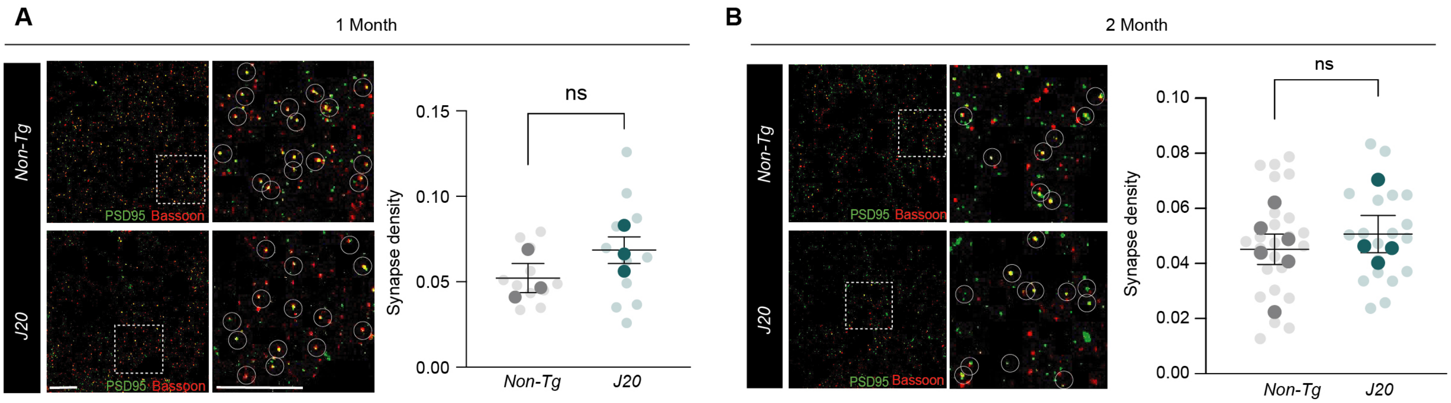


**Figure S7. *J20* mice have no change in synapse density at 1 and 2 months.**
(A-B) *J20* and Non-Tg cortical synapse density shows no significant difference at 1 month or 2 months of age. Synapse density was defined as number of Bassoon and PSD95 colocalized puncta normalized to area. All data are mean ± SEM with n=3-6 mice with Student's t-test for (A, B). Scale bar is 10 µm and 5 µm.

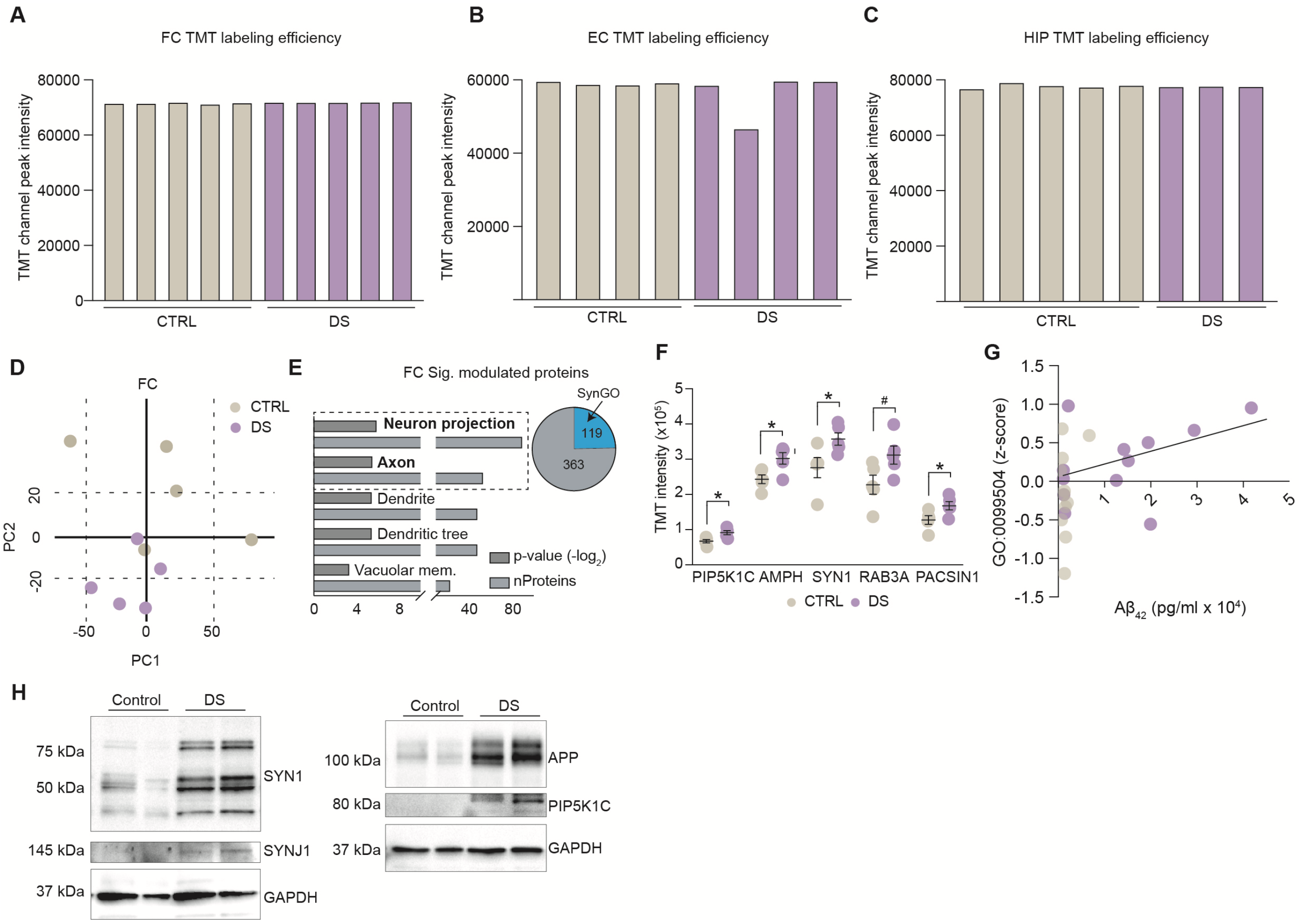


**Figure S8. TMT-MS labeling efficiency and biochemical validation of presynaptic protein accumulation in DS and CTRL brains.**

(A-C) TMT reporter ion intensities from the 10-plex TMT-MS experiments comparing DS and CTRL frontal cortex (FC), entorhinal cortex (EC), and hippocampus (HIP) extracts demonstrates equal labeling across all experiments.

(D) PCA analysis from the FC TMT experiment shows that biological replicates from DS and CTRL groups clusters together respectively.

(E) GO:CC analysis showing the cellular components overrepresented in the group of proteins with significantly increased abundance in the FC of DS compared to CTRL. Pie chart inset showing that about one-fourth of this protein pool are synapse associated.

(F) SV protein abundance (z-score value of proteins from GO:0099504) from all DS and CTRL brain region extracts plotted against respective $A\beta_{42}$ levels for each individual. DS patients show a positive correlation between $A\beta_{42}$ load and abundance of SV proteins.

(G) Representative WB analysis of FC extracts shows that DS patients have elevated presynaptic protein abundance (e.g., APP, PIP5K1C, SYN1, SYNJ1) compared to CTRL.
(H) TMT intensities for select "Presynaptic" proteins. All data are mean ± SEM for DS and CTRL human brains with FC (n = 5), EC (n = 4-5), and HIP (n = 3-5). # = p value = .05, * = p value < .05 by Student's t-test for (H).


1. N. R. Rao, J. N. Savas, Levetiracetam Treatment Normalizes Levels of Presynaptic Endocytosis Machinery and Restores Nonamyloidogenic APP Processing in App Knock-in Mice. *J Proteome Res* **20**, 3580-3589 (2021).
2. T. J. Hark, N. R. Rao, C. Castillon, T. Basta, S. Smukowski, H. Bao, A. Upadhyay, E. Bomba-Warczak, T. Nomura, E. T. O'Toole, G. P. Morgan, L. Ali, T. Saito, C. Guillermier, T. C. Saido, M. L. Steinhauser, M. H. B. Stowell, E. R. Chapman, A.

Contractor, J. N. Savas, Pulse-Chase Proteomics of the App Knockin Mouse Models of Alzheimer's Disease Reveals that Synaptic Dysfunction Originates in Presynaptic Terminals. *Cell Syst* **12**, 141-158 e149 (2021).
3. A. Upadhyay, D. Chhangani, N. R. Rao, J. Kofler, R. Vassar, D. E. Rincon-Limas, J. N. Savas, Amyloid fibril proteomics of AD brains reveals modifiers of aggregation and toxicity. *Mol Neurodegener* **18**, 61 (2023).
4. R. K. Carlin, D. J. Grab, R. S. Cohen, P. Siekevitz, Isolation and characterization of postsynaptic densities from various brain regions: enrichment of different types of postsynaptic densities. *J Cell Biol* **86**, 831-845 (1980).
5. Y. Z. Wang, C. C. M. Castillon, K. K. Gebis, E. T. Bartom, A. d'Azzo, A. Contractor, J. N. Savas, Notch receptor-ligand binding facilitates extracellular vesicle-mediated neuron-to-neuron communication. *Cell Rep* **43**, 113680 (2024).
6. T. L. Young-Pearse, J. Bai, R. Chang, J. B. Zheng, J. J. LoTurco, D. J. Selkoe, A critical function for beta-amyloid precursor protein in neuronal migration revealed by in utero RNA interference. *J Neurosci* **27**, 14459-14469 (2007).
7. N. R. Rao, A. Upadhyay, J. N. Savas, Derailed protein turnover in the aging mammalian brain. *Mol Syst Biol*, (2024).
8. J. N. Savas, Y. Z. Wang, L. A. DeNardo, S. Martinez-Bartolome, D. B. McClatchy, T. J. Hark, N. F. Shanks, K. A. Cozzolino, M. Lavallee-Adam, S. N. Smukowski, S. K. Park, J. W. Kelly, E. H. Koo, T. Nakagawa, E. Masliah, A. Ghosh, J. R. Yates, 3rd, Amyloid Accumulation Drives Proteome-wide Alterations in Mouse Models of Alzheimer's Disease-like Pathology. *Cell Rep* **21**, 2614-2627 (2017).
9. J. N. Savas, B. H. Toyama, T. Xu, J. R. Yates, 3rd, M. W. Hetzer, Extremely long-lived nuclear pore proteins in the rat brain. *Science* **335**, 942 (2012).
10. F. Koopmans, P. van Nierop, M. Andres-Alonso, A. Byrnes, T. Cijsouw, M. P. Coba, L. N. Cornelisse, R. J. Farrell, H. L. Goldschmidt, D. P. Howrigan, N. K. Hussain, C. Imig, A. P. H. de Jong, H. Jung, M. Kohansalnodehi, B. Kramarz, N. Lipstein, R. C. Lovering, H. MacGillavry, V. Mariano, H. Mi, M. Ninov, D. Osumi-Sutherland, R. Pielot, K. H. Smalla, H. Tang, K. Tashman, R. F. G. Toonen, C. Verpelli, R. Reig-Viader, K. Watanabe, J. van Weering, T. Achsel, G. Ashrafi, N. Asi, T. C. Brown, P. De Camilli, M. Feuermann, R. E. Foulger, P. Gaudet, A. Joglekar, A. Kanellopoulos, R. Malenka, R. A. Nicoll, C. Pulido, J. de Juan-Sanz, M. Sheng, T. C. Sudhof, H. U. Tilgner, C. Bagni, A. Bayes, T. Biederer, N. Brose, J. J. E. Chua, D. C. Dieterich, E. D. Gundelfinger, C. Hoogenraad, R. L. Huganir, R. Jahn, P. S. Kaeser, E. Kim, M. R. Kreutz, P. S. McPherson, B. M. Neale, V. O'Connor, D. Posthuma, T. A. Ryan, C. Sala, G. Feng, S. E. Hyman, P. D. Thomas, A. B. Smit, M. Verhage, SynGO: An Evidence-Based, Expert-Curated Knowledge Base for the Synapse. *Neuron* **103**, 217-234 e214 (2019).
